# A spatial atlas of colorectal cancer reveals the influence of stromal niches on tumour differentiation

**DOI:** 10.64898/2025.12.09.693138

**Authors:** Andrew D. Pattison, Rebekah M. Engel, Wing Hei Chan, Spencer Greatorex, David Nickless, Liam Skinner, Camilla Cohen, Julie Hickey, Christine Georges, Anne L. Fletcher, Paul J. McMurrick, Thierry Jardé, Lochlan Fennell, Helen E. Abud

## Abstract

Colorectal cancer is the third most common cancer worldwide and the second leading cause of cancer-related mortality. Tumour architecture is spatially heterogeneous, ranging from the necrotic core to the invasive front, accompanied by diverse stromal and immune responses that influence tumour progression and treatment outcomes. To explore the spatial organisation of the tumour microenvironment, we profiled 1000 genes in 846,469 cells in 23 normal and late-stage colorectal tumour samples. We identified nine distinct spatial niches based on their cellular composition. We show that lymphoid aggregates enriched for *CCR7*/*SELL*^+^ CD4 T cells displayed heightened interferon signalling, more proliferating B cells and gene expression changes indicative of an adaptive immune response. We defined granulocyte-rich regions are concomitant with inflammation-mediated stromal reprogramming and increased tumour stemness. Finally, we uncovered the impact of the tumour microenvironment by distinguishing gene expression programs that were intrinsic to cancerous epithelial cells from those mediated by niche-specific changes.

**Key points:**

1. We present a spatial atlas of colorectal cancer and normal colon alongside an R/shiny interface to interactively explore this data: https://abud-apps.abud-lab-spatial.cloud.edu.au/shiny/cosmxos/. We analysed nine CosMx slides and provide QC metrics, identify strengths and weaknesses and provide suggestions for analysis of imaging-based spatial data.
2. We identify and characterise nine distinct spatial niches, each with their unique gene expression profile.
3. We observe granulocyte infiltration associated with spatially controlled TNF and IL1 signalling from granulocytes and myeloid cells. This signalling associates with a reduction in pro-differentiation CXCL14⁺ fibroblasts and an increase in MMP⁺ tissue-remodelling fibroblasts, ultimately reprogramming tumours toward a more foetal/stem-like (progenitor) state. We suggest *IL24* as a potential regulatory factor of this response.
4. Granulocyte chemoattractants were consistently expressed at higher levels by both cancerous epithelial cells and the tumour microenvironment (TME) relative to normal colon.
5. We identify colonic LAs associated with T cell infiltration, including a population of *CCR7*/*SELL*^+^ CD4 T cells linked to significant reprogramming of B cells. Furthermore, we show that granulocyte-driven fibroblast and myeloid reprogramming is reversed within LAs.
6. We show that mutated epithelial cells within tumours are uniquely marked by an *FXYD5*^+^ *PIGR*^low^ expression profile, allowing for tumour vs normal epithelial cell differential gene expression analysis. This distinction clarifies cancerous tumour-intrinsic expression changes versus those promoted by the TME.

## Introduction

Colorectal cancer (CRC) is a leading cause of cancer-related death worldwide with projections for the year 2040 estimating 3.2 million new cases and 1.6 million deaths annually^1^. CRC tumours are heterogeneous and are composed of genetically mutated, cancerous epithelial cells (ECs) that are supported or impeded by a variety of stromal and immune cell types within the tumour microenvironment (TME)^2–5^. CRC is commonly understood to arise through the accumulation of genetic driver mutations that contribute to tumour progression through the dysregulation of well-known signalling pathways including WNT, MAPK and TGF-β^6,7^. Additionally, changes in the immune environment have been linked to prognosis^8^. Emerging evidence now suggests that the innate plasticity of colonic ECs can also be exploited to promote tumour progression, metastasis, and therapy resistance^9–14^. Although these mechanisms are still being explored, it is clear that tumours are influenced by extrinsic signals within the TME^15^.

The advent of single-cell RNA sequencing (scRNA-seq) technology has provided unprecedented insights into the cellular heterogeneity of CRC. These methods have enabled the identification of diverse cell subpopulations with distinct molecular profiles and dysregulated signalling pathways that contribute to immune evasion and treatment resistance^15–22^. While many intratumoural interactions between cell types have been characterised at the single-cell level^16,17^, the spatial complexity of these interactions remains incompletely understood. Recently developed spatial transcriptomics methods now provide the opportunity to more comprehensively describe tumour tissue structure, combining single cell gene expression with the positional information of cells^23–26^.

Here, we utilise the Nanostring CosMx Spatial Molecular Imaging platform (CosMx)^23^ to generate a spatial transcriptomic atlas of normal human colon and treatment-naive late-stage CRC at single-cell resolution. We reveal tumour-specific tissue structures and explore spatial relationships between immune, stromal and ECs. By leveraging public scRNA-seq data in combination with our spatial data, we define how the tissue is reorganised and reprogrammed to support CRC using endogenous signalling pathways. We find that granulocyte infiltration into tumours consistently results in innate inflammation, pro-tumorigenic myeloid cell reprogramming and the depletion of pro-differentiation *CXCL14*^+^ fibroblasts. This allows tumour ECs to maintain an undifferentiated/progenitor epithelial phenotype. We additionally identify a *CCR7*/*SELL*^+^ CD4 T cell population highly prevalent within gut-associated lymphoid tissue (GALT) that is consistently associated with extensive B cell reprogramming, allowing us to identify key genes associated with adaptive immunity in the colon. Finally, we define a tumour-specific gene signature and use it to separate gene expression changes within ECs that are due to signals from the TME and those that are tumour-intrinsic.

Unravelling the intricacies of the spatial regulation of cell states within CRC provides new insights into tumour heterogeneity, while also defining spatially-mediated interactions between tumours and the TME that may be targeted for more effective therapeutic strategies.

## Results

### A spatial atlas of normal human colonic mucosa and late-stage CRC

To identify the spatial gene expression changes in advanced CRC, we generated spatially resolved single cell transcriptomic data from stage III and IV cancer patients, comprising 23 samples (15 primary tumours and 8 normal colon, Table S1) run across nine slides of the CosMx platform^23^ (Figure 1a) in multiple runs. Whole exome sequencing performed on adjacent primary tissue or organoids generated from this tissue confirmed the tumour specimens contained key driver mutations typically found in CRC^27^ (Table S1). Protein immunofluorescence (IF) markers-facilitated cell segmentation and ∼1,000 genes in 846,469 cells from 352 fields of view (FOVs) (Figure S1) were visualised at single-cell resolution.

**Figure 1.**
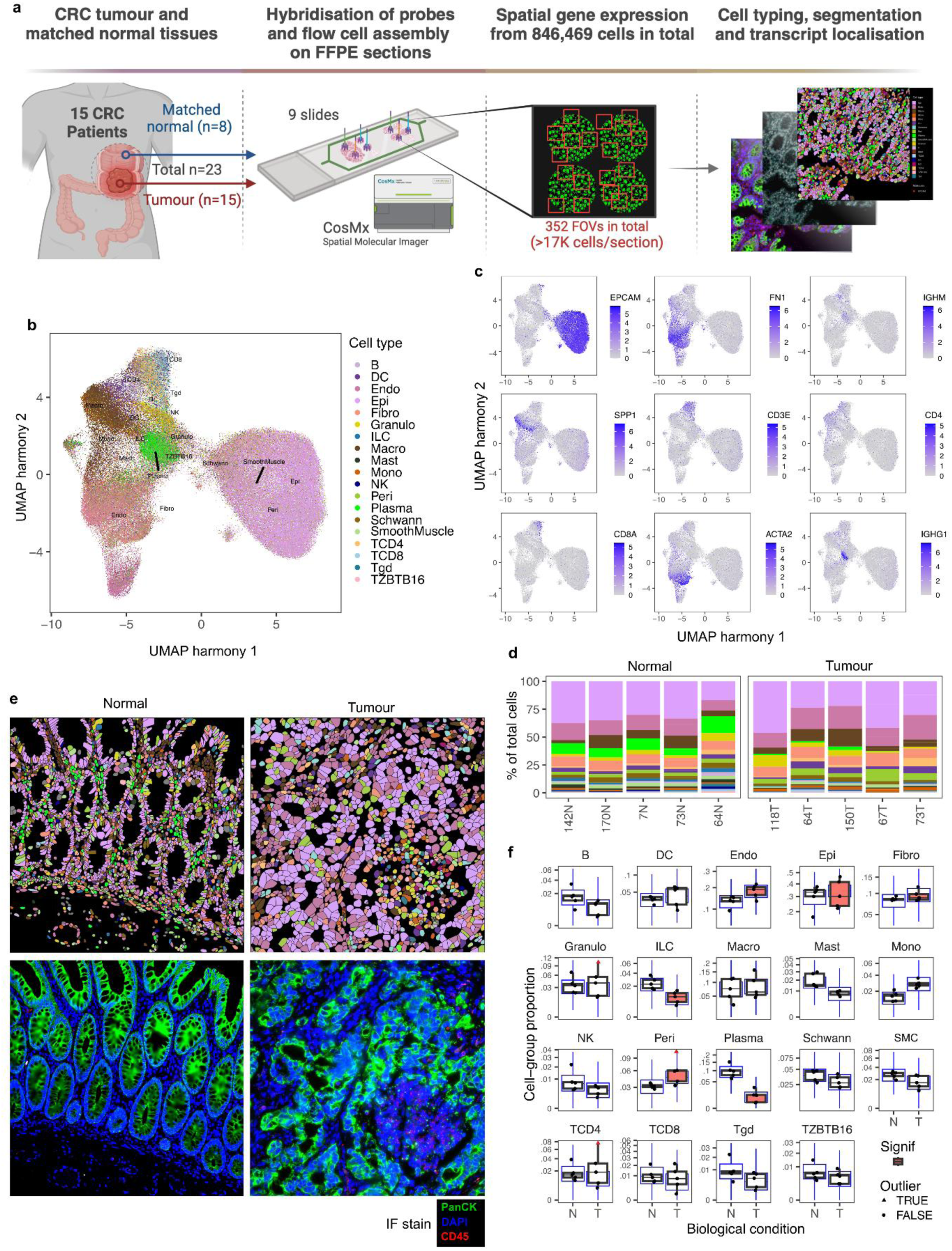
Annotation and composition of high-quality CosMx V1 cells. **a.** Schematic of the CosMx study design. **b.** UMAP of harmony integrated high-quality cells. Cell annotations provided by singleR using the Pelka single cell dataset as a reference. **c.** Canonical marker genes of known cell types. *EPCAM* marks ECs, *FN1* marks fibroblasts, *IGHM* marks IGM producing B cells, *SPP1* marks tumour-associated macrophages/monocytes, *CD3E* marks T cells, T cells can be split by *CD4* and *CD8*, *ACAT2* marks smooth muscle cells and *IGHG1* marks IGG B cells. **d.** The cellular composition of each sample split by tumour and normal. **e.** Representative images of SingleR cell type annotation overlaid onto a CosMx FOV from sample 73 normal (top) and tumour (bottom) and matching composite IF marker images (right). IF stains: green-PanCK, blue-DAPI, red-CD45. **f.** Changes in cell type proportions (DA) between tumour and normal samples as calculated by sccomp, red boxes indicate a significant association of sample type with cell composition, the blue boxplots represent a posterior predictive check.

To filter the data, CosMx-specific quality control was performed (see methods for full details). A median silhouette score (as employed by Stuart *et al*.^28^) was used to determine how well annotated cell types clustered together within a sample (Figure S2) and identify comparatively low-quality samples. Interestingly, the silhouette score was higher in well-differentiated tumours (Figure S3a) and was positively correlated with probe count (Pearson’s r = 0.69, p <0.01, Figure S3b), highlighting the importance of segmentation and RNA abundance for accurate cell type identification from spatial imaging technologies. Representative haematoxylin and eosin (H&E) images of well and poorly differentiated tumours are depicted in Figures S3c and S3e respectively. Of the 846,469 starting cells, 390,985 were annotated as high-quality. *NEAT1*, *MALAT1*, and negative probes (random probes that do not match a gene and control for background binding) were removed from analysis as their expression was variable and did not represent useful biological information (see discussion for more detail). This resulted in an average of 227 total and 107 unique probes detected per cell in Version 1 (V1) probe set and 339 total and 147 unique probes detected per cell in Version 2 (V2) probe set. We analysed each revision of the panel separately unless otherwise stated, but focussed on the V1 dataset as it contained more high-quality samples with a more diverse cellular composition.

To annotate the cells, we used scRNA-Seq data from mismatch repair deficient/proficient (MMRp/d) tumours and matched normal tissues from Pelka *et al*.^16^ (referred to as the MMR atlas) as a reference for automated and unbiased cell type annotation^29^. The mid-level annotations from the MMR atlas were selected as they were the highest resolution cell types that our CosMx data could reasonably be used to define, given the roughly 10 times lower per-cell read counts compared to standard scRNA-Seq. Consistent with cell type annotation, UMAP dimensionality reduction showed distinct clusters of epithelial, immune, and stromal populations (Figure 1b). These populations were additionally marked by the immunofluorescence for protein markers PanCK (epithelial), CD45/CD3 (immune/T cells) and CD68 (myeloid) populations (Figure S4).

The cellular composition and spatial distribution within the normal mucosa were largely consistent across samples (Figure 1d). These samples were obtained from different regions within the colon from patients with a variety of clinical characteristics including sex and age (Table S1). The normal mucosa was typified by a monolayer of columnar epithelium spatially organised into ordered colonic crypts, flanked by stromal and immune populations (Figure 1e left). In contrast, the architecture and cellular composition of tumours was highly disordered and variable, with immune and stromal populations dispersed throughout the lesions (Figure 1e right). Consistent with the changes in tissue organisation, the relative proportions of each cell type varied between tumour and normal samples (Figure 1f). In tumour samples we observed a significant reduction in the relative proportion of plasma cells and immature lymphoid cells (ILCs) and increases in the proportions of fibroblasts, pericytes, monocytes and macrophages (sccomp^30^ FDR < 0.05). Similar clustering and changes in cell type composition were also observed in the V2 probeset (Figure S5).

### Identification of a hallmark CRC transcriptional signature

To establish baseline differences between tumour and normal samples, pseudobulk differential gene expression (DGE) was performed. ECs from all tumour and normal samples were compared for both the V1 (Figure 2a) and V2 (Figure 2b) probesets. Pseudobulk EC counts from donor matched tumour and normal samples from the MMR atlas were also compared as a known truth (Figure 2c). Of note, *KRT23*, *IFITM1* and *IFITM3* were consistently increased in tumours, and *PIGR* and *FCGBP* were consistently decreased (FDR < 0.05, Tables S4/5/6). Reassuringly, the log_2_ fold changes (log_2_ FC) of significant genes (at FDR < 0.05) were highly correlated with those obtained when comparing donor matched tumour and normal samples from the MMR atlas scRNA-Seq dataset (Pearson’s r = 0.91, Figure 2d).

**Figure 2.**
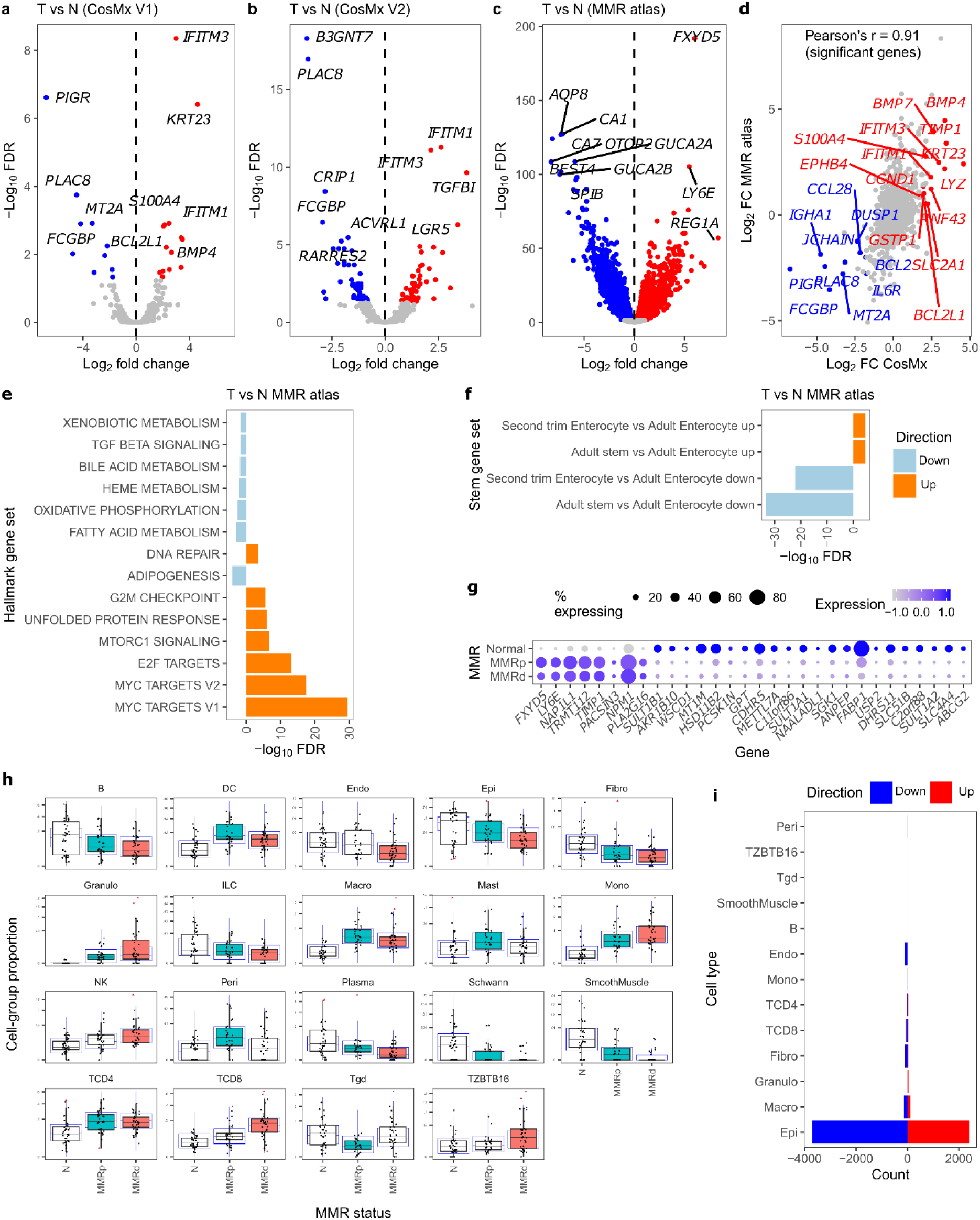
Comparing ECs from MMRp/MMRd tumours and normal tissue. **a**. Volcano plot of DE genes from a tumour vs normal contrast from the V1 CosMx probeset, the V2 probeset **(b)** and tumour vs matched adjacent normal in the MMR atlas (**c**). **d**. Correlation between significantly DE genes from the V1 CosMx probeset and the MMR atlas. **e**. Enrichment of hallmark gene sets and (**f**) foetal intestinal cell (second trim enterocyte) and adult intestinal stem cell gene signatures in tumour ECs in the MMR atlas. **g**. Dot plot of the top 30 genes from the gene sets used in **f** (ranked by FDR) in ECs split by MMR status. **h**. Changes in cell type proportions in samples split by MMR status as calculated by sccomp, colour indicates a significant association of sample type with cell composition, the blue boxplots represent a posterior predictive check. **i**. The number of DE genes in each cell type when comparing MMRd vs MMRp tumours.

Using a pseudobulk approach to compare tumour and normal samples highlighted additional tumour-specific gene expression changes to those specifically identified by the consensus non-negative matrix factorisation (NMF) method used by Pelka *et al*^16^. Key changes included large and highly consistent increases in the epithelial expression of *FXYD5, LY6E* and *REG1A* in tumours (Figure 2c, S12). Unfortunately, these genes were not represented in the CosMx panel. Limma-camera^31^ gene set enrichment analysis (GSEA) of hallmark gene sets enriched in tumour cells in the MMR atlas was also performed, showing expected tumour hallmarks, with increases in cell cycling most prominent (Figure 2e). Pseudobulk contrasts were also performed for all 18 cell types from V1 (Table S4), V2 (Table S5), the MMR atlas tumour vs normal (Figure S6, Table S6) and MMRp tumours vs MMRd tumours (Figure S7, Table S7), showing largely expected DE changes and providing useful reference lists of DE genes across multiple cell types from multiple conditions.

### CRC tumours revert to a progenitor, foetal-like state

In healthy colonic mucosa, the epithelium is renewed via LGR5^+^ stem cells that reside at the base of crypts^32^, but recent single cell studies have identified other transient stem cell populations marked by a foetal signature (*CLU*, *ANXA1*, *PROX1*^9^) that appear in response to injury^33,34^ and are implicated in CRC tumour recurrence^13,15,35^. To interrogate stem cell signatures in CRC tumours, we again ran GSEA on the MMR atlas (tumour vs normal EC contrast) using four custom foetal and stem cell gene sets defined from the human Gut Cell Atlas (https://www.gutcellatlas.org/, see methods for gene set definition). Tumours displayed a stem/progenitor CRC phenotype with a reduction in differentiation markers, more closely resembling both stem cells and foetal enterocytes (Figure 2f). Colonic stem cell and regenerative markers *LGR5*, *YAP1*, *CLU* and *ANXA1* were higher in tumour ECs compared to normal in the matched MMR atlas (Table S6, FDR <0.01), with these trends also reflected in the CosMx data (Table S4/S5). Notably, *FXYD5* and *LY6E* that were higher in our initial tumour vs normal analysis (Figure 2c) were also strong foetal signature genes (Table S8) and were consistently more highly expressed regardless of MMR status (Figure 2g). Indeed, changes in cell type proportions and gene expression were largely consistent between MMRp/d tumours with some exceptions; e.g. increased CD8 T cell infiltration into MMRd tumours (Figure 2h) and MMR-associated changes in tumour gene expression (Figure 2i).

*FXYD5* (also known as dysadherin) was clearly the most consistently upregulated gene in MMR atlas tumours regardless of MMR status (Figure 2c, Table S7), but was not discussed by Pelka *et al*.^16^ and is surprisingly not well studied in CRC. In normal colon ECs marked by *EPCAM* expression (Figure S8a), FXYD5 was notably absent in normal ECs compared to other cell types (Figure S8b, left panel) but present in tumour ECs. *FXYD5* is known to promote EMT through the downregulation of E-cadherin (*CDH1*) in multiple other cancer types^36–38^. Concordantly, *CDH1* expression was also consistently lower in tumours (FDR <0.01, log_2_FC = -1.2). *FXYD5* has also been suggested to promote growth and metastasis in epithelial ovarian cancer by enhancing transforming growth factor-β (TGF-β) signalling^39^. While T vs N GSEA showed that TGF-β signalling was slightly lower in tumour cells relative to normal (FDR = 0.02, Figure 2e, Table S12), *TGFBI* was the third most consistently increased DE gene (Figure S8c, log_2_ FC = 5.4, FDR = 1.1×10^-76^), suggesting a specific TGF-β signalling response despite a reduction in the expression of some pathway members overall.

In the context of the colon, the function of the second most consistently upregulated gene *LY6E* is also relatively poorly studied. It has, however, previously been proposed as a prognostic marker, with knockdown reducing CRC proliferation *in vitro*^40^. Interestingly, *LY6E* was expressed in most cell types, but was most restricted in normal ECs (Figure S8d), suggesting tight control of this gene under normal conditions. *PIGR* was another consistently downregulated gene that has been reported to be involved in immune signalling and negatively correlated with outcome in CRC^41^. *PIGR* transports polymeric immunoglobulins (IGs) to the intestinal lumen where they can be secreted^42^. *PIGR* mutations have been found to be under positive selection in colonic crypts of patients with inflammatory bowel disease^43^, but have not been reported in cancer. *PIGR* was also massively downregulated in foetal compared to adult colonocytes from the human gut atlas (log_2_FC = -13.26, FDR < 0.01, Table S8), suggesting this change is also symptomatic of a reversion to a more foetal-like state. Interestingly, a dramatic reduction in the expression of markers of the *BEST4*^+^ mature absorptive epithelial subtype^44,45^, was observed in tumours including down-regulation of *OTOP2*, *CA7*, *GUCA2A*, *GUCA2B*, and *SPIB*, indicating that differentiated *BEST4^+^* cells are largely absent in CRC. Overall, this analysis revealed that regardless of the molecular subtype, CRC tumours exhibit signatures representative of a progenitor foetal state, with *FXYD5*, *LY6E* upregulated and *PIGR* expression reduced.

### Identification of spatial niches within CRC

To identify discrete spatial microenvironments, samples were partitioned into nine cellular neighbourhoods or ‘niches’ using the Seurat BuildNicheAssay workflow^23,46,47^. This method counted the 50 closest spatial neighbours (k = 50) of each cell and then partitioned cells into niches based on patterns of similar neighbouring cell composition using K-means clustering. We chose nine niches as this was consistent with previous cellular neighbourhood identification in CRC by Schürch *et al*.^48^. Niches were then manually annotated based on their cellular makeup (Figure 3a). We observed similar niches to Schürch *et al*.^48^ including epithelial mass (tumour), granulocyte-rich, LAs (follicle), vasculature (vascularised smooth muscle) and stroma (immune infiltrated stroma). Niche compositions per-FOV were largely consistent within samples (Figure S9), with some exceptions for the tumours, likely due to immune responses or necrosis (discussed further below). The niche composition of each sample was again more consistent for normal colon as compared to tumour (Figure 3b). Examples of niche mapping onto FOVs are shown in Figure 3c (ii) for normal and tumour (iv). Significant differences in niche composition were observed between tumour and normal samples, with tumours having greater proportions of epithelial and stromal rich niches, while normal samples mostly consisted of ‘normal-like’ niches, which reflected the structure of colonic crypts (Figure 3d). Of note, immune rich LAs were identified in both tumour and normal samples, while granulocyte-rich niches were exclusively observed in tumours.

**Figure 3.**
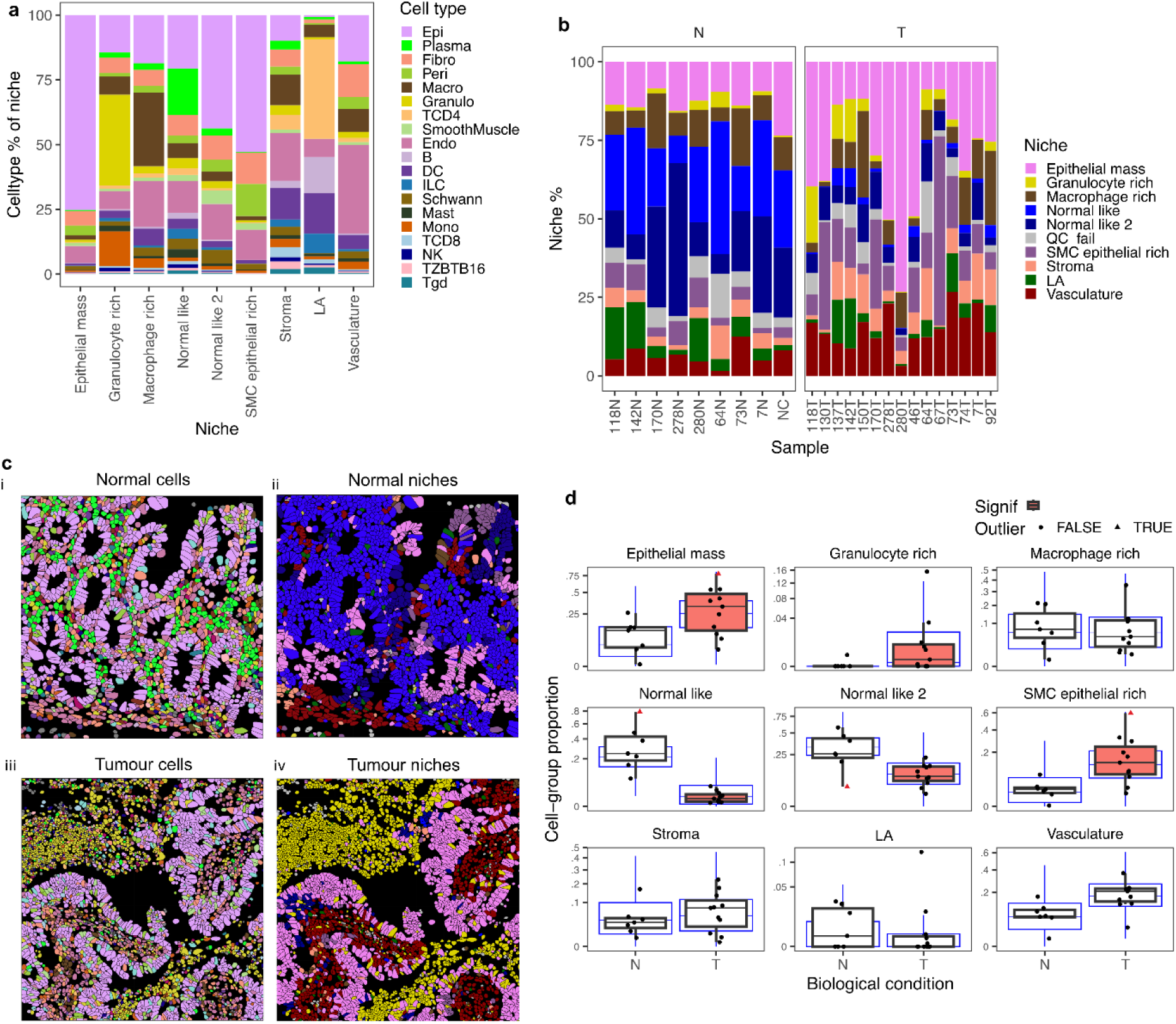
Categorisation of cells into 9 spatial niches. **a.** Cell type compositions of identified spatial niches. **b.** Niche composition of tumour and normal samples including a repeated normal negative control (NC). **c.** Representative images of SingleR cell type annotation overlaid onto a CosMx FOV from sample 142N FOV 4 (i) and 118T FOV 8 (iii) and their respective niche annotations (i and iv). **d.** Changes in niche proportions (DA) between tumour and normal samples.

### LAs contain *IL7R*^+^ *SELL*^+^ CD4 T cells associated with B cell activation

The immune system plays a major role in modulating outcomes in CRC with the distinct roles of immune components that react to tumour neoantigens increasingly emerging^3,49^. Currently, cytokine therapies, checkpoint inhibitors, peptide vaccines, CAR-T cells and targeting of the complement system are all being explored as potential immunomodulatory treatment options for CRC^50^. The niche that is best known to be associated with tumour outcome and immune checkpoint blockade (ICB) response is the tertiary lymphoid structure (TLS)^51–53^. This is an inflammation-induced immunological niche characterised by separated T and B cell aggregates, resembling lymph nodes or splenic white pulp^54,55^. While we identified similar structures in our CosMx data, we did not have sufficient per-cell gene expression counts to separate TLSs from gut-associated lymphoid tissues (GALTs) present in normal colon^56^ and so we defined this population more broadly as the LA niche. These structures were chiefly composed of CD4 T cells, B cells, DCs and ECs, with the largest proportion made up by CD4 T cells (Figure 3a). Pseudobulk contrasts of cells in this niche against all other niches revealed high expression of the DC markers *CCL19*, *LAMP3* and *CCR7* by fibroblasts, ECs, CD4 T cells and DCs (Figure 4a). DCs expressing these genes are often defined as mature DCs enriched in immunoregulatory molecules (mregDCs) that may be associated with improved T cell function and checkpoint immunotherapy response^57^.

**Figure 4.**
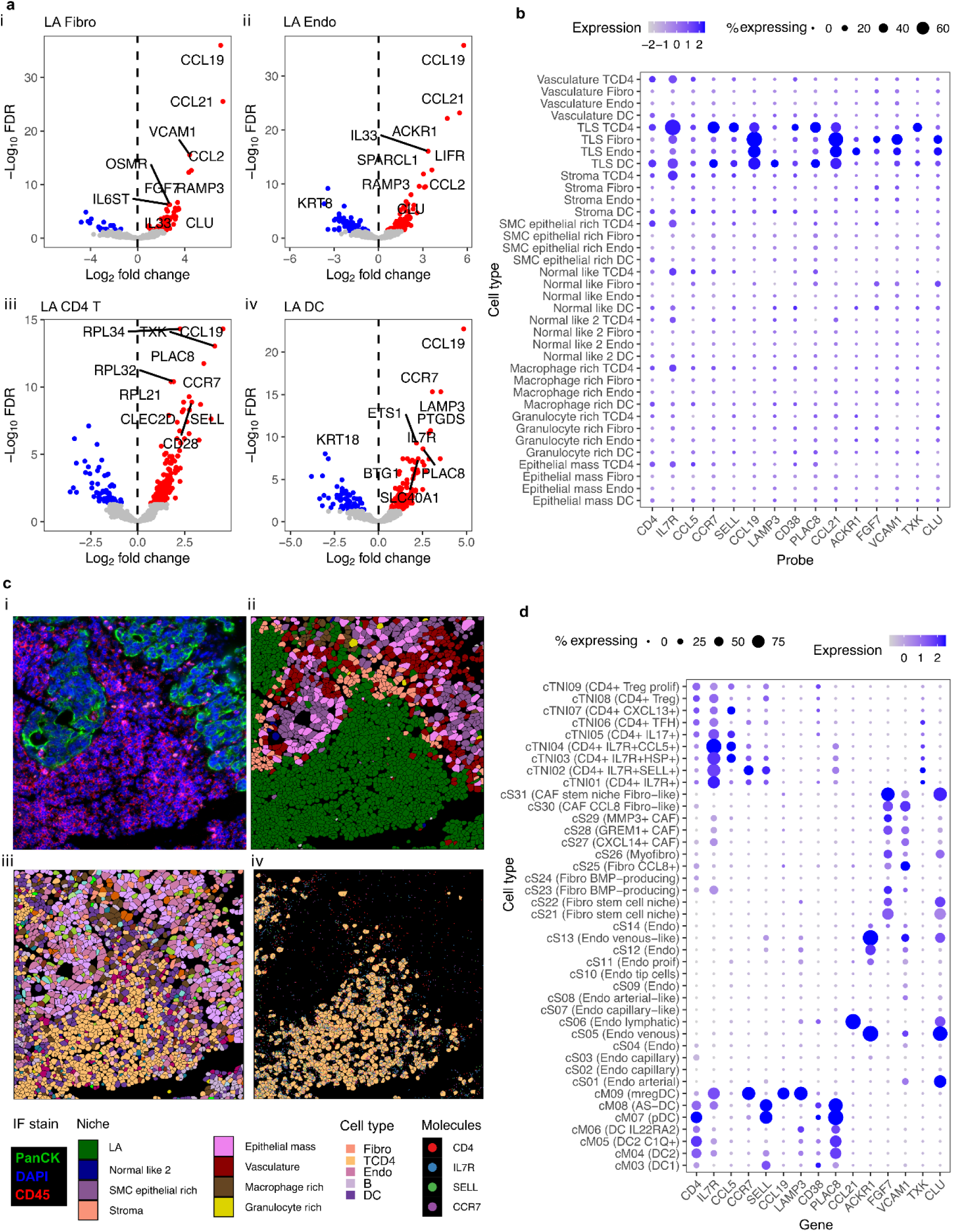
CosMx characterisation of LAs reveals a novel *IL7R*/*SELL* CD4 T cell subset. **a.** Volcano plots of the top DE genes comparing LA to all other niches for FBs (i), ECs (ii), CD4 T cells (iii) and DCs (iv). **b.** Dot plot of the expression of some key genes from **a** acoss all niches. **c.** Example of a LA niche in CosMx sample tumour 73 FOV 39. IF staining (i), niche annotation (ii), cell type composition (iii) and key LA CD4 T cell probes *CD4*, *IL7R SELL* and *CCR7* (iv). **d.** The same genes from b in FB, EC, CD4 T (iii) and DC subtypes from Pelka *et al*^16^.

The T cell chemoattractant *CXCL9* was also significantly upregulated in macrophages within the LAs, with a concurrent downregulation of *SPP1*, which conversely, was strongly expressed in the granulocyte-rich niche (Table S9). Clonal tumour mutation burden (TMB) in combination with *CXCL9/13* expression have also been suggested as the strongest predictors of ICB response^58^. Interestingly, the DC marker *CCR7* was also higher in LA-CD4 T cells, which also had increased expression of *SELL* and *IL7R* (Figure 4b), indicating a potentially LA-specific CD4 T subtype (Figure 4c). To confirm that this gene expression pattern was not due to inaccurate cell segmentation, the same genes were plotted from the highest resolution MMR atlas cell types, confirming that these genes were indeed expressed in a subset of CD4 T cells (Figure 4d).

Based on the expression profile within CD4 T cells in the CosMx LAs, we hypothesised that the cTNI02 (*IL7R*^+^*SELL*^+^ CD4) cell type identified in the MMR atlas^16^ reflected CD4 T cells within tumour LAs. Splitting the MMR atlas into tumours with the highest numbers of these cells (‘LA-high’) and performing pseudobulk DE analysis (Figure S10, Table S10) revealed a striking increase in both the number and activation state of B cells (Figure 5a/b). There was an increase in IGD^+^IgM^+^ GC−like and CD40^+^ GC−like B cell populations (Figure 5a) and significant gene expression changes in B cells generally. Indeed, despite splitting the dataset on a CD4 T cell subset, B and tumour cells showed a higher number of significant changes between sample groups (Figure 5b), suggesting *IL7R*^+^*SELL*^+^ CD4 T cells may indeed be key components of LAs.

**Figure 5.**
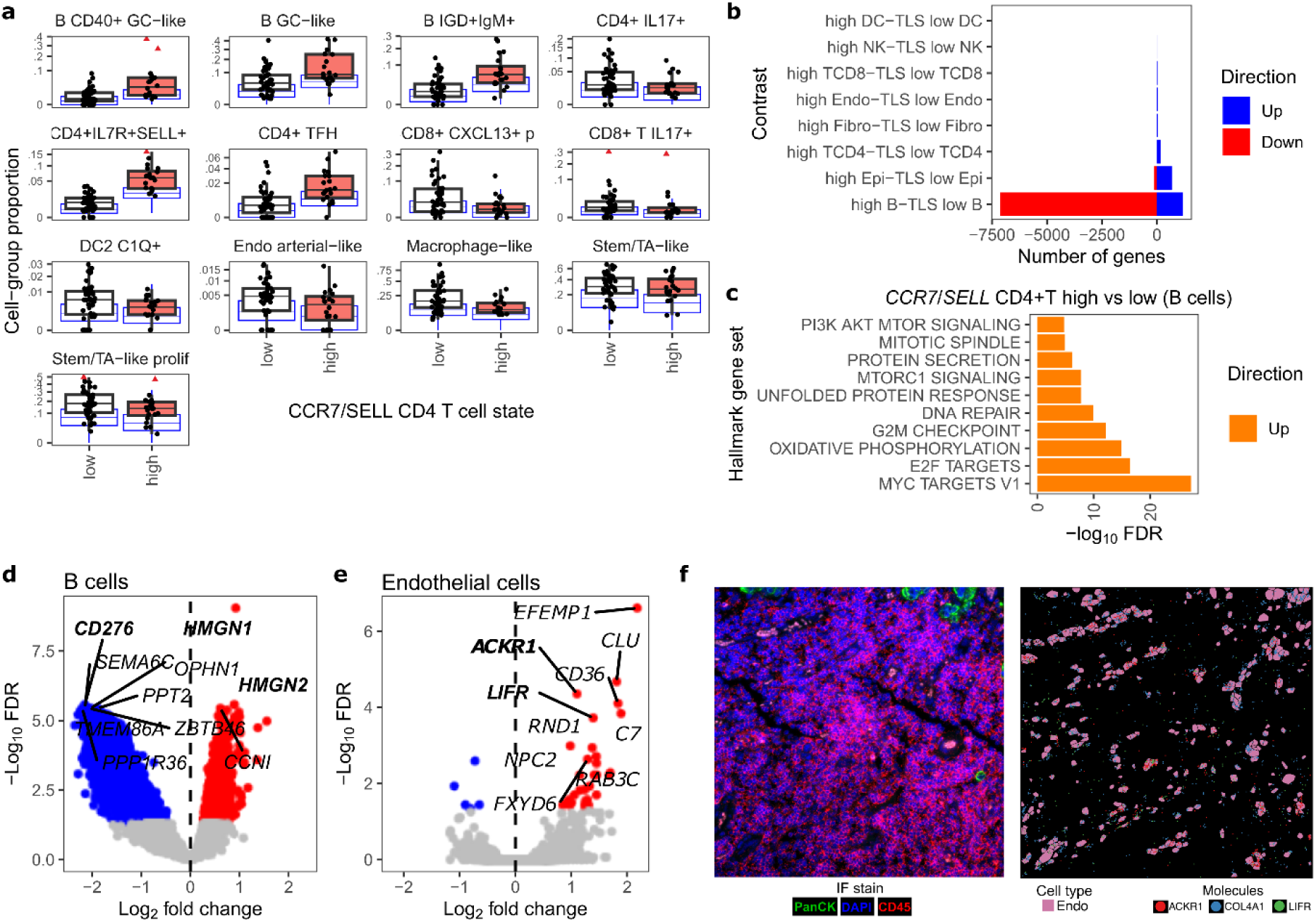
Partitioning MMR atlas tumours into LA high/low subsets based on CosMx niches reveals striking changes in B cell infiltration and EC gene expression. **a.** Changes in MMR atlas low-level cell type proportions with high/low LA infiltration. **b.** The number of DE genes per mid-level cell type in LA high vs low tumours. **c.** Hallmark gene sets enriched in LA high B cells. **d.** Volcano plot of DE genes between LA high low samples for B cells and ECs (**e**). **f.** IF staining of a LA (left), and the ECs in this FOV (right). *COL4A1* is shown as a marker of ECs and *ACKR1* and *LIFR* are shown as they are DE in LA high samples in the MMR atlas.

LA low vs high GSEA analysis further showed a broad increase in B cell metabolism and cell cycling (Figure 5c) including the downregulation of the immune checkpoint *CD276*^59^ and increased expression of *HMGN1* and *HMGN2* (Figure 5d), which may be associated with naive B cell recruitment and transcriptional activation^60^. The expression of *ACKR1 and LIFR* was also significantly higher in LA-high endothelial cells (Figure 5e), which was strikingly consistent with the CosMx V1 LA niche (Table S9, Figure 5f). Increased expression of *ACKR1* suggested organisation of a chemokine gradient to direct cellular influx^61^, perhaps in response to increased IFN signalling in these regions as predicted by GSEA (Table S10). Taken together, these results describe in clearer detail the cellular composition and expression profile of this key immunogenic region.

### Granulocyte chemoattraction is strongly driven by myeloid cells and CAFs

In addition to LAs, CosMx spatial niche identification also revealed striking granulocyte-rich regions which were poorly infiltrated with other immune cell types (Figure 3c/6a). These regions were tumour-specific (Figure 3d) and surrounded isolated aggregates of likely necrotic ECs^16^. Consistent with the cellular composition of these regions, increased expression of granulocyte chemoattractants was observed in fibroblast and myeloid populations from both the MMR atlas tumours (Figure S11a) and the CosMx granulocyte-rich niche (Figure S11b). *CXCL8 is* the most potent neutrophil chemoattractant, and is opposed by *CXCL12*^62^. Consistently, *CXCL8* had the greatest increase in expression in MMR atlas tumour fibroblasts when compared to normal (log_2_ FC = 6.7, FDR = 1.16 x 10^-26^, Figure S11c), accompanied by a decrease in *CXCL12* (log_2_ FC = -2.79, FDR = 7 x 10^-16^, Figure S11d). Interestingly, in the CosMx dataset neutrophil chemoattractant expression was also most reduced in LA niches (Table S9, Figure S11b).

### Granulocyte infiltration is associated with pro-tumorigenic myeloid cells and CAFs

We observed striking examples of concentrated granulocyte infiltration in two CosMx FOVs (Figure 6a), however, a relatively low number of granulocyte-rich FOVs were present in the CosMx dataset overall (Figure S9) and analysis was limited to the 1,000 genes in the CosMx panel. Therefore, to investigate the gene expression changes associated with granulocyte infiltration in more samples, we revisited the MMR atlas dataset which has confirmed granulocyte infiltration, identified as likely neutrophils, from imaging validation^16^. We split the tumours in this dataset into high and low infiltration groups and compared them, once again also modelling sex and MMR status as factors that were not of interest. However, we note that despite being generally increased in tumours, granulocyte infiltration was higher in MMRd tumours compared to MMRp tumours, with 1.9% vs 0.4% of detected cells annotated as granulocytes respectively, with this trend approaching significance (p = 0.086, two-sided Wilcoxon rank-sum test). Consistent with the CosMx granulocyte-rich niche, *SPP1* was again significantly higher in the monocyte population of the granulocyte-high cohort (Figure 6b) and approaching significance in the macrophage population (FDR = 0.08, Table S11). Additionally, while no metastases were analysed in this study, *MRC1* was also higher in myeloid cells from the granulocyte-rich MMR samples (log_2_ FC = 2.5, FDR <0.01, Table S11). *MRC1* has been associated with the TME of CRC liver metastases^63^. Macrophages also showed a decrease in *DNASE1L3* (log_2_ FC = -4.8, FDR <0.01), which has a role in the degradation of neutrophil extracellular traps (NETs)^64^ that cause vascular occlusion, possibly counteracting a potentially tumour growth limiting effect of granulocyte infiltration.

**Figure 6.**
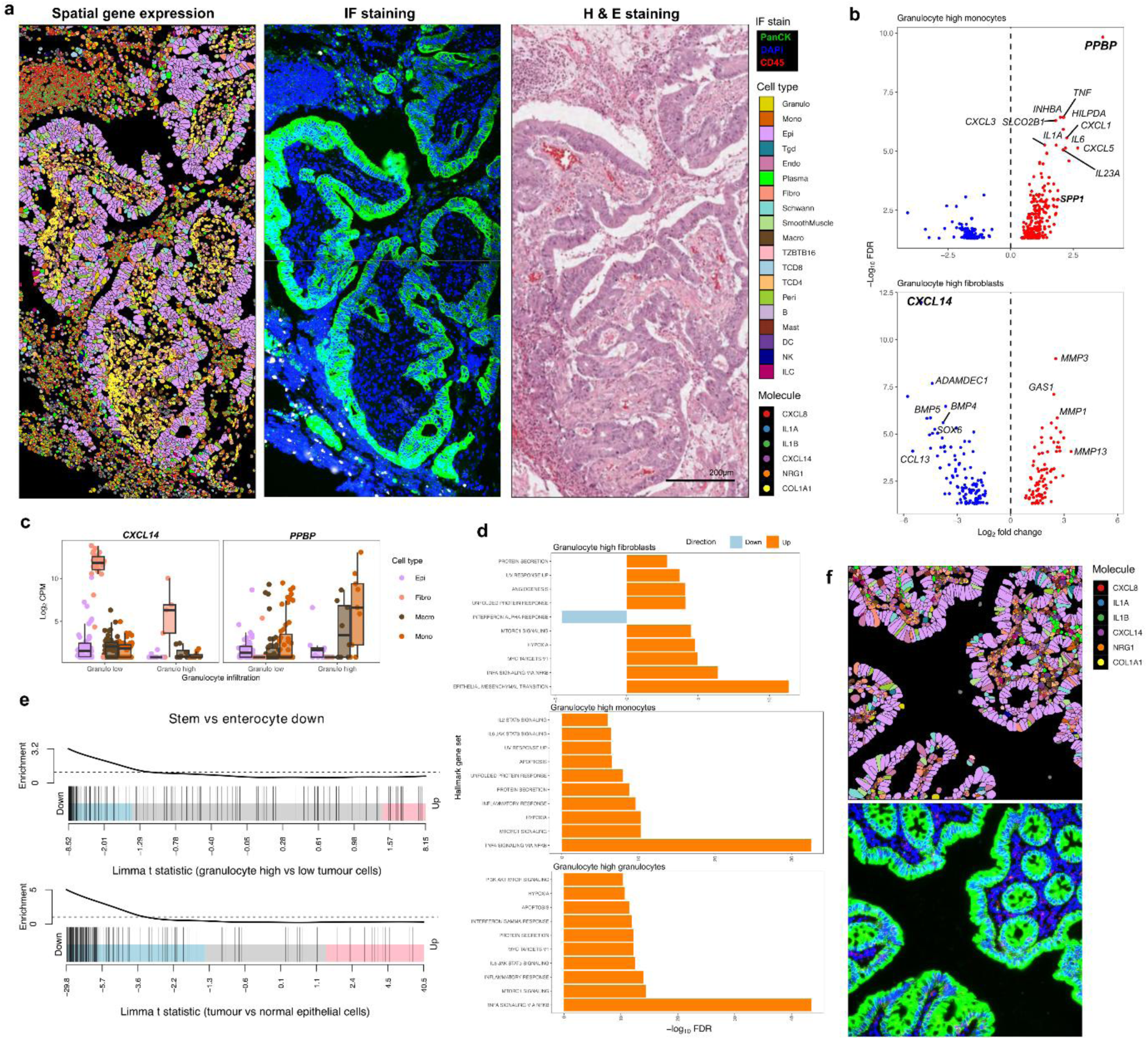
Neutrophil infiltration decreases fibroblast mediated tumour cell differentiation. **a** Neutrophils infiltrating a likely necrotic region of tumour 118T and with high expression of *CXCL8*, *IL1A* and *IL1B*. Fibroblasts marked by *COL1A1* lack expression of *CXCL14* and *NRG1*. Matched IF composite and H & E staining is shown to the right. **b.** Volcano plots showing the top 20 DE genes (by FDR) when comparing cell types with the greatest number of gene expression changes in tumours with low vs high granulocyte infiltration. Both patient sex and tumour MMR status were accounted for before contrasting these cell types. Single cell data is from the Pelka MMR atlas. **c.** Top Limma-camera enriched hallmark gene sets in non-tumour cell types present in **b**. **d.** Barcode plots showing the downregulation of genes in the stem vs enterocyte down gene set derived from the gut cell atlas. Downregulation of genes associated with enterocyte differentiation is enriched in granulocyte high tumours (top, FDR = 3.56 x10^-13^) and in tumour cells compared to normal epithelial tissue (bottom, FDR = 1.65 x10^-34^). **e.** Boxplots highlighting decreased *CXL14* expression in granulocyte high fibroblasts (left) and increased *PPBP* in granulocyte high monocytes and macrophages (right). **f.** Example of *CXCL14* and *NRG1* expression by normal fibroblasts in sample 142N.

Fibroblast activation protein alpha (*FAP*), which was not present in the CosMx panel, was also higher in fibroblasts and pericytes in the granulocyte-high MMR atlas cohort (Table S11). A recent report showed reduced response to anti-PD-L1 therapy in patients with high *SPP1*/*FAP* expression and a lack of T cell infiltration into these tissues^17^. Interestingly, *IL24*, which is implicated in wound repair and the inhibition of proliferation and migration of keratinocytes^65^ was rarely detected in normal tissue fibroblasts. In contrast, *IL24* was elevated in tumours (Figure S12a, Table S6). This difference was again more pronounced in granulocyte-rich samples (Figure S12b, Table S11). *IL24* has been positively associated with CRC survival^66^ and an oncolytic virus containing *IL24* has also been shown to inhibit tumour growth in CRC models^67^. We therefore hypothesise that *IL24* may help to prevent uncontrolled growth of fibroblasts, and potentially ECs as well, in response to inflammation associated with granulocyte infiltration, however experimentally validating this is beyond the scope of this study.

The clearest changes in the granulocyte-rich samples were in monocytes, with an increase of *PPBP* expression (Figure 6c), which has been reported to potently attract and activate neutrophils^68^. Also significantly upregulated were the innate inflammatory markers *IL1A* and *IL1B*^69^ (Table S11). This increase in innate inflammation was accompanied by a significant decrease in the expression of the T cell chemoattractants *CXCL9/10/11*. GSEA of the same contrast also revealed a large increase in TNF-ɑ signalling in monocytes (Figure 6d). This increase in TNF-ɑ and wound healing may be broadly supportive of tumour growth, indeed an anti-TNF-ɑ mAb has been shown to limit tumour growth in an orthotopic CRC mouse model^70^.

### Granulocyte infiltration is associated with fibroblast-mediated suppression of epithelial cell differentiation

Tumours with granulocyte infiltration showed a striking decrease in the expression of genes associated with the differentiation of mature enterocytes compared to colonic stem cell markers (Figure 6d top). This same reduction was also seen when comparing tumour and normal samples (Figure 6d bottom), which is consistent with the undifferentiated phenotype of tumour ECs, but also the minimal granulocyte infiltration within normal colon noted previously. Pelka *et al*.^16^ described a *FRZB, BMP, CXCL14* expressing subset of fibroblasts that directs the differentiation of normal colon ECs. We also observed this *CXCL14^+^* subset expressing *NRG1* in some populations of fibroblasts in our CosMx data (Figure 6f), but not in the granulocyte-rich niche (Figure 6a, Table S9). Expression of *CXCL14* was also significantly reduced in granulocyte-rich tumours in the MMR atlas (Figures 6b/e), but not significantly DE between tumour and normal fibroblasts (Table S6), further supporting the notion that this reduction is driven by granulocyte infiltration.

In addition to signals targeting ECs, fibroblasts in granulocyte-rich tumours from the MMR atlas also upregulated *CXCL5*, *MMP1*, *MMP3* and *IL6.* This is consistent with inflammation due to infection and tissue remodelling. *CXCL14*, *BMP4/5* and *FRZB*, were expressed at lower levels and this may drive EC differentiation^16^. It is possible that bacterial infiltration within the colon triggered inflammation, however, there was a decreased response to IFN-ɑ (Figure 6d) and lower expression of the antimicrobial *CCL13* (Figure 6b, Table S11), which would suggest the response was more limited to tumour necrosis. Taken together, these results support a model where inflammation associated with tumour necrosis results in T cell infiltration, supports tissue remodelling and helps to maintain a foetal/stem like state in tumours.

### A *FXYD5^+^ PIGR* low expression profile defines cancerous epithelial cells

We next sought to decipher which expression signatures in tumour ECs were tumour-intrinsic compared to those that are instigated by niche environments within the TME. We exploited our findings that *FXYD5* expression is high in tumour ECs and PIGR expression is high in normal ECs to define different EC populations. Once again utilising single cell data from the MMR atlas we defined tumour ECs as *FXYD5^+^ PIGR* low and normal ECs as *FXYD5^-^ PIGR* high. Of the 87,329 starting ECs, neither gene was detected in 19,856 ECs and both genes were detected in 6,533 ECs. Both classes were marked as ambiguous and removed from further analysis as they could not be confidently assigned to either tumour or normal. The remaining 60,940 cells were then partitioned into predicted tumour or normal cells (Figure 7a). To confirm normal cells could be effectively separated from tumour samples, inferCNV was run on sample 157 as a representative example (Figure 7b). Samples from this donor clustered by tumour status and only predicted tumour cells contained chromosomal abnormalities (Figure 7c) and a significant number of more normal ECs were detected in this sample. When comparing the proportions of EC subtypes, sccomp showed that most tumour cells appear undifferentiated and lack the hallmarks of differentiated cell types as expected. (Figure 7d).

**Figure 7.**
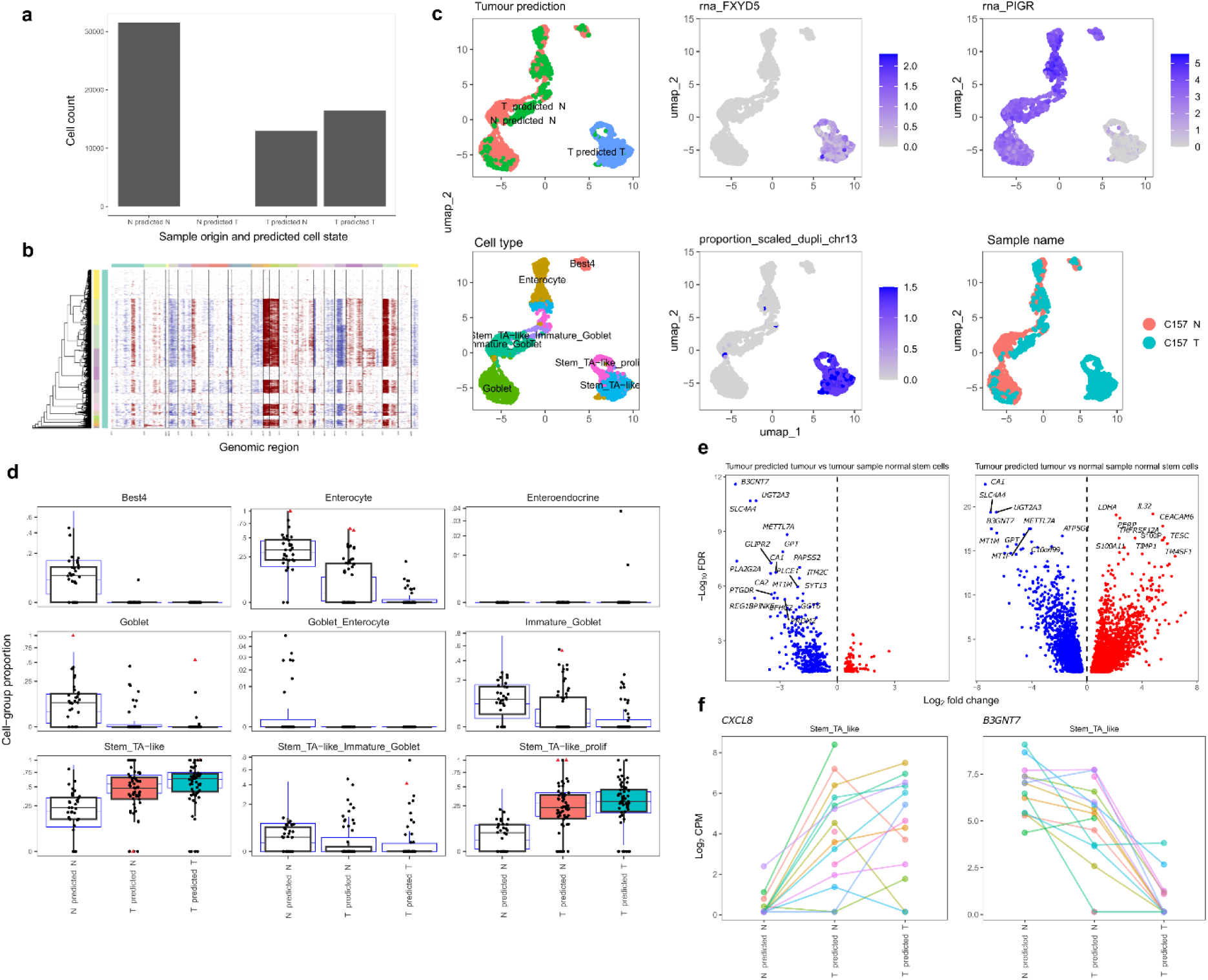
A tumour-specific gene expression signature reveals even non-tumour cells within the tumour niche are stem-like. **a.** The numbers of ECs in the MMR atlas assigned to either tumour or normal cell types within each sample type by loss of *PIGR* and activation of *FXYD5* expression. **b** InferCNV copy number changes in donor 157 from the MMR atlas confirms many cells in the tumour sample appear not to have chromosomal alterations. **c**. UMAPs of ECs from donor 157 showing that tumour cells largely cluster separately from normal sample cells and cells predicted as normal in the tumour sample. These cells were effectively split by loss of *PIGR* and activation of *FXYD5* expression. Most tumour cells were annotated as stem-like and had chromosomal instability as shown by the change in Chr13. **d**. Boxplots from sccomp comparing EC type proportions in normal samples, predicted normal cells from tumour samples and predicted tumour cells from tumour samples. **e**. Volcano plot of top DE genes comparing predicted tumour stem/TA like cells from tumours with predicted normal stem/TA like cells from tumours (left) and predicted tumour stem/TA like cells from tumours with predicted normal stem/TA like cells from matched normal samples (right). **f**. *CXCL8* is generally increased in tumour samples regardless of if the cell is of tumour origin, whereas *B3GNT7* expression is decreased substantially in tumour cells specifically.

Separating mutated tumour ECs from tumour-derived normal ECs allowed us to define a transcriptional signature of these populations. We focussed on the ‘stem/TA-like’ cells as the majority of tumour cells fell into this category, making it the most relevant comparison (Figure 7e). These analyses revealed further tumour-intrinsic gene expression differences. We noted a large decrease in *B3GNT7* expression specifically in tumour ECs alongside upregulation of *IL32* and *CEACAM6*. Interestingly, we did not detect any significant changes between tumour-derived normal ECs and tumour ECs in molecules that attract neutrophils (Table S13). This included *CXCL8*, which is highly upregulated when comparing tumour ECs and normal sample-derived normal ECs (Figure 7f). This suggested that neutrophil chemoattraction is a general response to EC proliferation (and likely necrosis) rather than being tumour-specific.

GSEA results from the within tumour stem cell comparison mostly showed gene expression changes related to cell cycling (Table S14). These changes were also present in the tumour vs normal EC stem cell populations, but were overshadowed by changes that were likely driven by the TME, including a clear shift to hypoxic metabolism, EMT, response to TNF-ɑ and response to IFN-ɑ/γ. Taken together, this data suggests that the overall profile of CRC ECs is comprised of intrinsic mutated cancerous tumour EC gene expression (such as increased proliferation), in combination with a normal EC-like response to the TME. We present a model of these combined processes in Figure 8.

**Figure 8.**
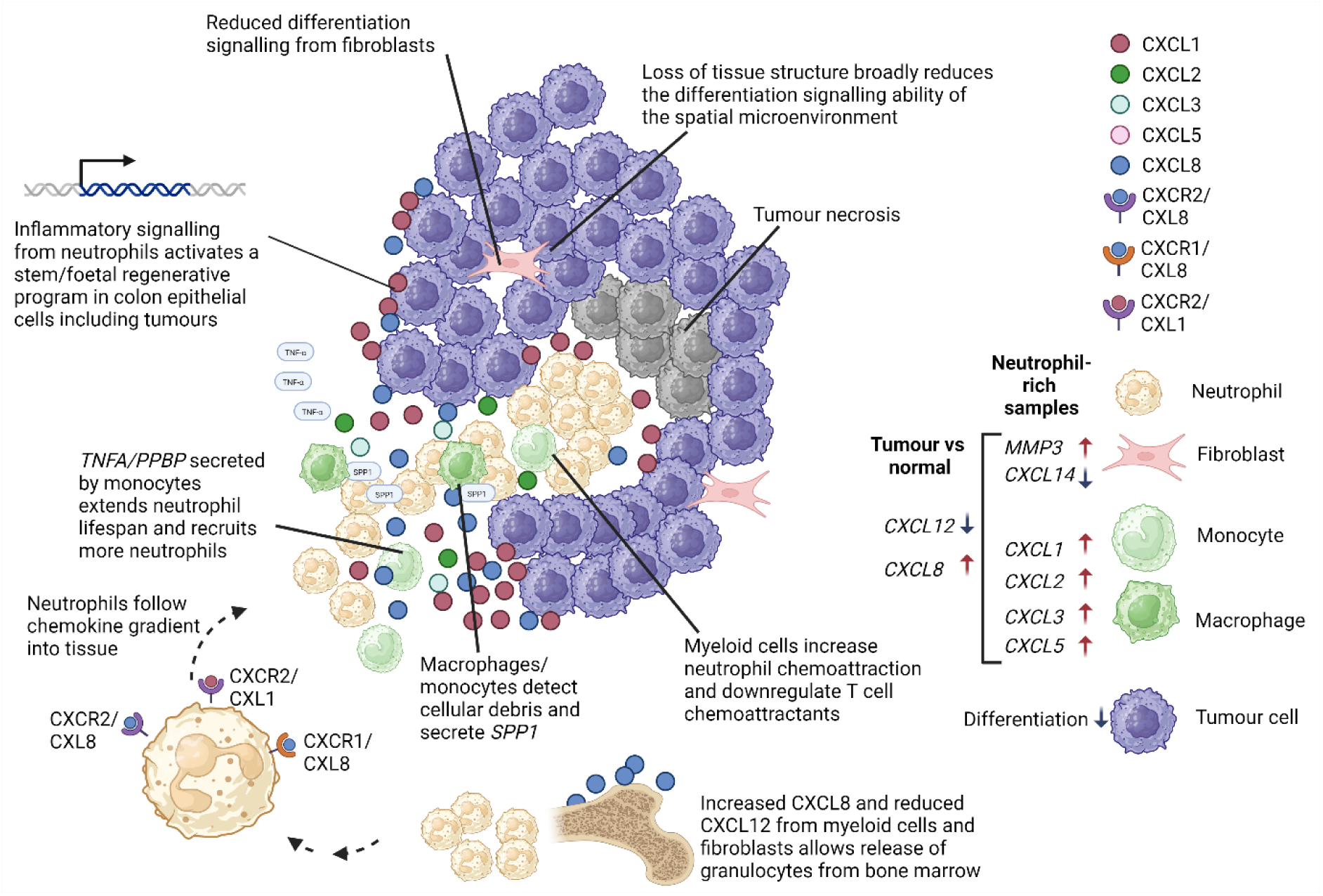
Proposed model of neutrophil-driven tumour stemness. Tumour necrosis leads to the infiltration of neutrophils into the tumour tissue. Inflammatory signalling including TNF-ɑ drives regenerative program in both the stroma a colonic epithelium that drives a stem/foetal program in colon epithelial cells (including tumours). Created in BioRender. Pattison, A. (2025) https://BioRender.com/s99p57z

## Discussion

The interaction between tumour cells and both immune and stromal cell types in the surrounding tissue microenvironment are key determinants of tumour progression and response to therapies^5,81–84^. Here, we used spatial transcriptomics to identify and characterise niches within primary colon tumours and normal colon to identify mechanisms that may support or inhibit tumour growth.

We show that primary colorectal tumours from treatment-naive patients with late-stage disease exhibit an undifferentiated foetal-like signature, and that this response is exacerbated by granulocyte infiltration. *FXYD5* and *LY6E* were identified as both key markers of this signature and definitive markers of tumour epithelial cells. *FXYD5* encodes dysadherin, a membrane-penetrating protein that can inactivate E-cadherin and is upregulated in many solid tumours and is associated with tumour progression^85–87^. Dysadherin knockout in mice decreases tumour incidence in both genetic (*Apc^Min/+^*) and chemically-induced models of CRC^86^. Functional studies suggest that dysadherin has a role in promoting tumour growth by impacting cell adhesion and promoting epithelial to mesenchymal transition (EMT). A monoclonal antibody has been developed against dysadherin for use in the diagnosis of pancreatic and lung cancer^88^, highlighting the potential for this to also be applied to CRC tumours. *LY6E* has been identified as a prognostic marker associated with worse overall survival in CRC, with knockdown of *LY6E* in cell lines observed to reduce cellular proliferation *in vitro*^40,89^. The striking consistency of increases in the expression of these genes in CRC paired with their association with foetal cell state put them forward as strong candidates for future investigation.

Spatial analyses conducted in this study provided further insight into the signals that augment foetal epithelial expression profiles associated with cellular plasticity during progression of CRC^9^. Our study defined distinct niches within tumours where changes in EC states were associated with profound alterations in stromal and immune cell populations. Inflammation, likely driven by tumour necrosis, creates a microenvironment that is granulocyte-rich and T cell depleted, and activates a regenerative response in stromal cells that enables and maintains dedifferentiation of ECs in tumours. Our data supports the concept of stromal expression of neutrophil chemoattractants such *CXCL8* and *CXCL5* and the neutrophil response/activators *SPP1/PPBP* by monocytes.

Epithelial and stromal cells in close proximity to granulocytes also displayed a clear shift in transcriptional profile. This dedifferentiated epithelial state polarised tumours to be much more foetal-like and is accompanied by a TNF-ɑ response alongside a specific and consistent loss of genes that mark the differentiated and short lived *BEST4*^+^ absorptive subtype. Notably, this included the polymeric immunoglobulin receptor (*PIGR*). Taken together, this model (Figure 8) shows that inflammation related to tumour necrosis generates a regenerative response in the TME which in turn activates a foetal stem-like phenotype, which enabling further proliferation of mutated ECs in tumours. Foetal reversion has also been associated with metastasis and chemoresistance^9,90^. Recent work has shown that it is possible to re-differentiate CRC cell lines in *vitro*^91^. Given the endogenous pathways that allow tumour dedifferentiation presented here, it may be possible to devise treatments that terminally differentiate cancerous ECs. In support of this concept, patients with RNA profiles suggesting a higher level of pro-differentiation CXCL4-expressing fibroblasts appear to have a better prognosis^92^.

The cellular composition and functional states within LAs were further characterised as part of this study. Aggregates of T and B cells described as TLSs are associated with a better prognosis in CRC and are proposed to support an environment that is immunoresponsive^54,93,94^. The precise function of cells within the LA remains to be defined and is confounded by the presence of GALTs, which are also present in normal colon and may have similar spatial and gene expression patterns^56^. Despite this limitation, our data provide insight into the spatial gene expression profiles of B and T cells present within these regions.

Application of the CosMx platform enabled unprecedented exploration of the CRC TME, however, there are still some limitations that may also apply to other imaging-based spatial technologies. As this atlas was compiled over multiple CosMx runs (Figure S17a), there were changes to probes (Table S15) and the resolution of the CosMx imager (Figure S17b), which made integration of data challenging. The high expression of lncRNAs such as *NEAT1* and *MALAT1* (Figure S17c) also necessitated the removal of these transcripts. *NEAT1* and *MALAT1* are also known to have an association with active chromatin^95^ rather than cell type. As may be expected given the limitations of optical density on a CosMx slide, cell size was generally correlated with the number detected probes (Figure S17d) and the number of cells of a given type that were detected (Figure S17e). This bias may explain the underrepresentation of some cell types including NK and T cells in CosMx compared to scRNA-Seq (Figure S18a/b).

Gene expression profiles detected in CosMx highly correlated with scRNAseq and V1 and V2 cell type proportions and probe detection were broadly consistent (Figure S18c, f). The CosMx data did not reach saturation, as an increase in the number of probes detected per cell continued to yield additional unique probes (Figure S19), indicating that further transcript detection is possible with increased probe capture. It is for these reasons that spatial predictions were not directly assigned to single cell data using tools such as CeLEry^96^, however, this may be possible with future improvements to technology or spatial analysis methodologies.

Negative probe expression appeared to be an issue on slides 4 and 5 (Figure S20a), with some of the highest detection of any probes (Figure S17c), however this issue was seemingly resolved in the commercial version (V2) of the CosMx protocol (Figure S21a). Across both runs ∼50% of the captured cells were discarded as there wasn’t enough transcript count to reliably determine the cell type of those cells (Figures S20/S21 b). Once confounding cells and probes were removed, median per-cell probe counts were ∼200 (Figures S20/S21c) and unique probe counts were ∼100 (Figures S20/S21d) depending on the sample. In both V1 and V2 some of the most highly expressed and highly variable genes were plasma cell genes (Figures S20/S21e), which are often overrepresented in the ambient RNA profile of scRNA-Seq samples. Indeed, the plasma cell markers *JCHAIN*, *IGHA1* and *IGKC* also seemed to be present in most other cell types (Figure S16a), suggesting that slide-based imaging data may not be immune from this problem. Combined these issues made it difficult to resolve clear clusters in both V1 (Figure S22) and V2 (Figure S23). Further increases in the number of detected probes and a reduction of background noise are required before the per-cell transcript quality of imaging-based spatial transcriptomic methods approaches that of scRNA-Seq.

This study contributes a powerful dataset that demonstrates the organisation of spatial niches in the normal colon compared to primary late-stage colorectal cancer. Our analysis revealed transcriptional programs that define tumour epithelial states associated with the spatial localisation and expression programs of surrounding stromal and immune cells. This spatial context of cells within the TME can either support or inhibit disease progression as well as determine tumour differentiation state. Delineating these cellular relationships and defining signals that promote changes in cellular state will provide critical insights into the mechanisms that drive aggressive cancer phenotypes to advance the development of effective therapies.

## Methods

### Patient tissue samples

This study was conducted in accordance with the Declaration of Helsinki, and the protocol was approved by the Cabrini Research Governance Office (CRGO 04-19-01-15 and 04-15-05-17) and the Monash University Human Research Ethics Committee (MUHREC ID 2518). Tissue specimens were obtained following surgery from treatment naïve patients at Cabrini Hospital, Malvern, Australia. All patients provided written informed consent. Stage III and IV primary colorectal tumours from 15 patients and normal colonic mucosal samples from 8 patients were included in the study. Clinical information, including patient and tumour characteristics, were obtained from the prospectively maintained, clinician-led Cabrini Monash University Department of Surgery colorectal neoplasia database (CMCND)^71^. Patient age at time of tissue collection ranged from young onset (27 years) through to older donors (88 years), with a median age of 77 years (Table S1). There was no attrition of subjects over the course of the study.

### Determination of oncogene mutations by whole exome sequencing

Organoids were grown, harvested and underwent WES sequencing as described previously^27^. Fastq files were aligned to the hg38 using the nf-core^72^ sarek pipeline^73^ (version 3.4.0). Somatic variants were identified with the SuperFreq pipeline^74^ (version 1.6) running under R (version 4.4.0), with 29 in-house normal organoids used as the panel of normals. Somatic variant annotation was performed using PCGR (version 2.0.1)^75^.

### Tissue microarray construction

Tissue microarrays were constructed at the Monash Histology Platform. 2 mm cores were dissected from formalin fixed paraffin embedded clinical samples using a Micro Tissue Arrayer, MTA-1 (Beecher Instruments, USA). Core placement was guided by the histopathological annotation of tissue blocks, excluding overly necrotic areas, as defined by an expert anatomical pathologist (Dr David Nickless). Donor cores were arranged into TMA recipient blocks and reannealed at 45 degrees for 3 hours. 4µm sections were collected onto slides. To aid in field of view (FOV) placement, the first and last slide in each series of sequential sections was stained with haematoxylin and eosin (H&E) and digitised at 40X magnification on the Aperio AT slide scanner. High resolution H&E images are available as supplemental materials (Figure S15).

### CosMx SMI sample preparation and experimental setup

The CosMx Human Universal Cell Characterization Panel was utilised to profile the expression of 1000 genes in normal colon and tumour tissue specimens^23^. Two versions of the 1000-plex gene panels were used across the dataset: Version 1 (V1) was employed under the Nanostring Technology Access Program (slides 1–5), while Version 2 (V2) was utilized at the Central Facility for Genomics, Griffith University (slides 6–9) (Table S15). Cell segmentation was performed using IF cell membrane and morphology marker proteins DAPI, cell segmentation marker CD298/B2M, PanCK and CD45 (all slides), CD3 (slides 1-3) and CD68 (slides 5-9). Transcript-specific probes are hybridized to the RNA molecules on FFPE slides, detected by the CosMx SMI, and cleaved to allow subsequent rounds of cyclical hybridizations and readouts. This allowed for gene identification, detection of the spatial location of RNA molecules and assignment cells.

### CosMx QC filtering and sample integration

Probe counts, probe spatial positions and cell segmentation information were exported as flat files (CSVs) from the Nanostring AtoMx platform. CosMx FOVs from the same sample/slide combination were first analysed individually for QC purposes. Probe counts from each slide were read into R (version 4.4.0) using the LoadNanostring function from Seurat (version 5.1.0)^46^. Negative and system control probes were removed along with the highly expressed lncRNAs *NEAT1* and *MALAT1.* Cells with less than 50 total detected probes were removed and any probes that were not subsequently detected in at least 50 cells were also excluded. For each cell, the median absolute deviation (MAD) was calculated for the total reads and total features detected. Cells with a MAD value > 3 for either of these metrics were also filtered out. While the doublet profile of CosMx is likely different to that of single cell, incorrect segmentation may result in incorrect assignment of transcripts to cells. Therefore, scDblFinder^76^ was run with default settings to remove the top 10% of cells that most resembled heterotypic doublets. Counts were scaled using the Seurat ScaleData, and we subsequently performed PCA, UMAP and clustering analysis using RunPCA, RunUMAP, FindNeighbors, FindClusters functions, respectively, with 50 PC dimensions used at the RunUMAP and FindNeighbors steps.

To determine the cell types present within each cluster, singleR (2.6.0) was run with default settings, using the pre-integration raw counts with the MMR atlas^16^ mid-level cell type annotations as the reference. Annotated cell types consisted of those derived from the myeloid lineage (dendritic cells - DC, monocytes - mono, macrophages - macro, granulocytes – granulo, and mast cells - mast), eight of lymphoid origin (CD4^+^ T-lymphocytes - TCD4, CD8^+^ T-lymphocytes - TCD8, B-lymphocytes - B, plasma cells - Plasma, natural killer cells - NK, immature lymphoid cells - ILC, gamma-delta T-lymphocytes - Tgd, and ZBTB16^+^ T-lymphocytes), five stromal populations (fibroblasts - fibro, pericytes - peri, endothelial cells - endo, schwann cells - schwann, and smooth muscle cells - smooth muscle) and Epithelial Cells - Epi. Due to lower per-cell transcript counts in CosMx data compared with 10x scRNA-Seq, the MMR atlas mid-level annotations were subsequently used for all downstream analyses. To verify our annotations, we looked at additional sources including the human primary cell atlas^77^ (Figures S13/S14, V1/V2), cell type marker genes (Figure 1c) and mean IF from protein markers recorded by the CosMx machine (Figure S4, V1), all of which largely concurred with MMR atlas mid-level annotations. The CosMx probes marking these cell types (ordered by P value) can be seen in Figure S16a (V1), with the full lists in Table S2 (V1) and Table S3 (V2). To evaluate the consistency of clustering and cell type annotations we used the ‘silhouette’ function from the ‘cluster’ R package^78^. Sample/slide combinations with visually confirmed acceptable clustering (median silhouette score > -0.025) were retained for further analysis.

Due to differences in probesets (Table S15) and protocols, samples from the Nanostring technology access program (TAP) (V1 probeset) and the commercial (V2 probeset) were analysed separately following QC. Starting with the raw gene expression values from the previously retained cells, log normalisation and variable gene selection were performed using the Seurat NormalizeData and FindVariableFeatures functions with default settings. The counts from the selected genes were then scaled and PCA analysis was performed. Harmony^79^ was used to integrate the data for clustering/visualisation based on the PCA reduction, mitigating the effects of the sample and slide batches. Further sub clustering of these populations did not separate the cells into identifiable subtypes (Figure S16b). Sub Clustering paired with the relatively low per-cell read counts (compared to scRNA-Seq) in this dataset suggested that it would be difficult to identify cell types at a higher resolution.

### Niche identification

The Seurat FindNeighbors function was run using the spatial coordinates of each cell to make a matrix of the 50 closest spatial neighbours. This matrix was then scaled and centred using the Seurat ScaleData function before k-means clustering with a k value of 9, consistent with a previous CRC imaging study^48^. Niches were then manually annotated based on their cell type composition.

### CosMx tumour vs normal

Cells were grouped based on sample, CosMx slide and cell type. Groups containing <15 cells were dropped. Probe counts from each group were then summed to form pseudobulk samples. Only one sample/slide combination was used to avoid repeated contrasts from the same sample on different slides disproportionately influencing DE results. Each cell type from tumour samples was then contrasted against its normal counterpart using limma-voom^31^. Slide level batch effects estimated and removed using limma duplicateCorrelation. Patient sex was mostly balanced across tumour and normal conditions so was not modelled to preserve degrees of freedom DE testing. Samples were not randomised across slides.

### MMR atlas tumour vs normal

Raw gene level counts from Pelka *et al*.^16^ were downloaded from the gene expression omnibus (GEO) accession: (GSE178341). To keep only high-quality cells and exclude doublets, the percentage of reads mapping to mitochondrial genes, the total read count and the number of genes detected were all required to have a MAD < 2 relative to other cells in the same sample. Consistent with the CosMx annotation, the mid-level annotations from the study were used as cell type identifiers. The counts for each cell type within each sample were summed (pseudobulked) and differential gene expression was performed between donor matched tumour and normal samples using limma-voom.

### Definition of foetal and stem cell gene sets

Raw gene level counts from cells from the Gut Cell Atlas^80^ were pseudobulked by author-defined sample and cell type annotation and contrasted using limma-voom. Stem cell gene sets were defined as those up (172 genes) or down (164 genes) at a FDR < 1×10^−10^ when comparing adult stem cells to adult enterocytes. Foetal gene sets were defined as those up (294 genes) or down (323 genes) at a FDR < 1×10^−20^ when comparing second trimester enterocytes to adult enterocytes. Gene sets are provided as a GMT: Gene Matrix Transposed file format (*.gmt) file in the supplemental material.

### Contrasting granulocyte and LA high and low MMR atlas samples

MMR atlas tumour samples were split into granulocyte high (>2 % of total cells per sample) and low (<2% of total cells per sample) groups based on a histogram of granulocyte infiltration, with nine high and 55 low samples identified. Similarly the MMR atlas was split into LA high (>1% *CD4 IL7R*^+^*SELL*^+^ of total cells per sample) and low (<1% *CD4 IL7R*^+^*SELL*^+^ of total cells per sample) resulting in 20 high and 40 low samples identified. Pseudobulk differential gene expression and limma-camera GSEA analyses were performed, with patient sex and MMR status modelled as variables not of interest focus on the effects of high granulocyte infiltration or LA presence respectively.

### Splitting tumour samples into tumour and normal subsets

ECs from matched tumour and normal samples from the MMR atlas were annotated as ‘predicted tumour’ if they had any detectable *FXYD5* and ‘predicted normal’ if normalised *PIGR* (seurat lognorm counts) were > 1. Cells that contained both markers or where neither marker was detected were dropped. While tumour vs normal DE analyses suggested the respective specificities of *FXYD5* and *PIGR*, we also ran inferCNV of the Trinity CTAT Project (https://github.com/broadinstitute/inferCNV) on sample 157 to as a measure of the CNV profile of the predicted tumour and normal populations. To assess changes in the epithelial population sccomp^30^ DA analysis was performed comparing predicted tumour and predicted normal from tumour samples against the predicted normal cells (all cells) from the normal samples with the previously encoded cell type annotations with the highest levels of specificity used. Pseudobulk DE analysis was next performed between the same groups as previously described.

## Supporting information

All supplemental figures

All supplemental tables

## Data and code availability

Spatial transcriptomics data generated using the CosMx SMI instrument is available at GEO: (GEO accession GSE303070). We have also developed an interactive app to explore this data: https://abud-apps.abud-lab-spatial.cloud.edu.au/shiny/cosmxos/. Scripts used to analyse these data are available at https://github.com/Abud-lab/cosmx-atlas.

## Author Contributions

HEA, RME, WHC, TJ, PJM, LF participated in study conceptualisation and study design. PJM, CG, RME, DN provided clinical expertise in selection of samples. RME, HA, LF, TJ, CC, JH, designed and prepared the tissue microarray. AP contributed bioinformatic analyses including the majority of figures. LF and LS assisted with data ingestion and formatting. LF contributed feedback and suggestions for bioinformatic analyses. AP, HEA, RME, WHC, TJ, PJM, LF, AF, SG performed analysis and interpretation of data. AP, RME, WHC, HEA, TJ, AF drafted the manuscript and all authors provided input on the final version.

## Acknowledgements

We thank the colorectal surgeons at Cabrini Health and the patients who participated in this study. The authors acknowledge the use of equipment and technical assistance of the Monash Histology platform, Department of Anatomy and Developmental Biology, Monash University. We thank the Monash Bioinformatics Platform, SMI Technology Access Program, and the Central Facility for Genomics, Griffith University for supporting work in this study. We thank Prof Wei Shi for providing helpful comments on the manuscript.

