## Supplementary material for "A spatial atlas of colorectal cancer reveals the influence of stromal niches on tumour differentiation": All supplemental figures

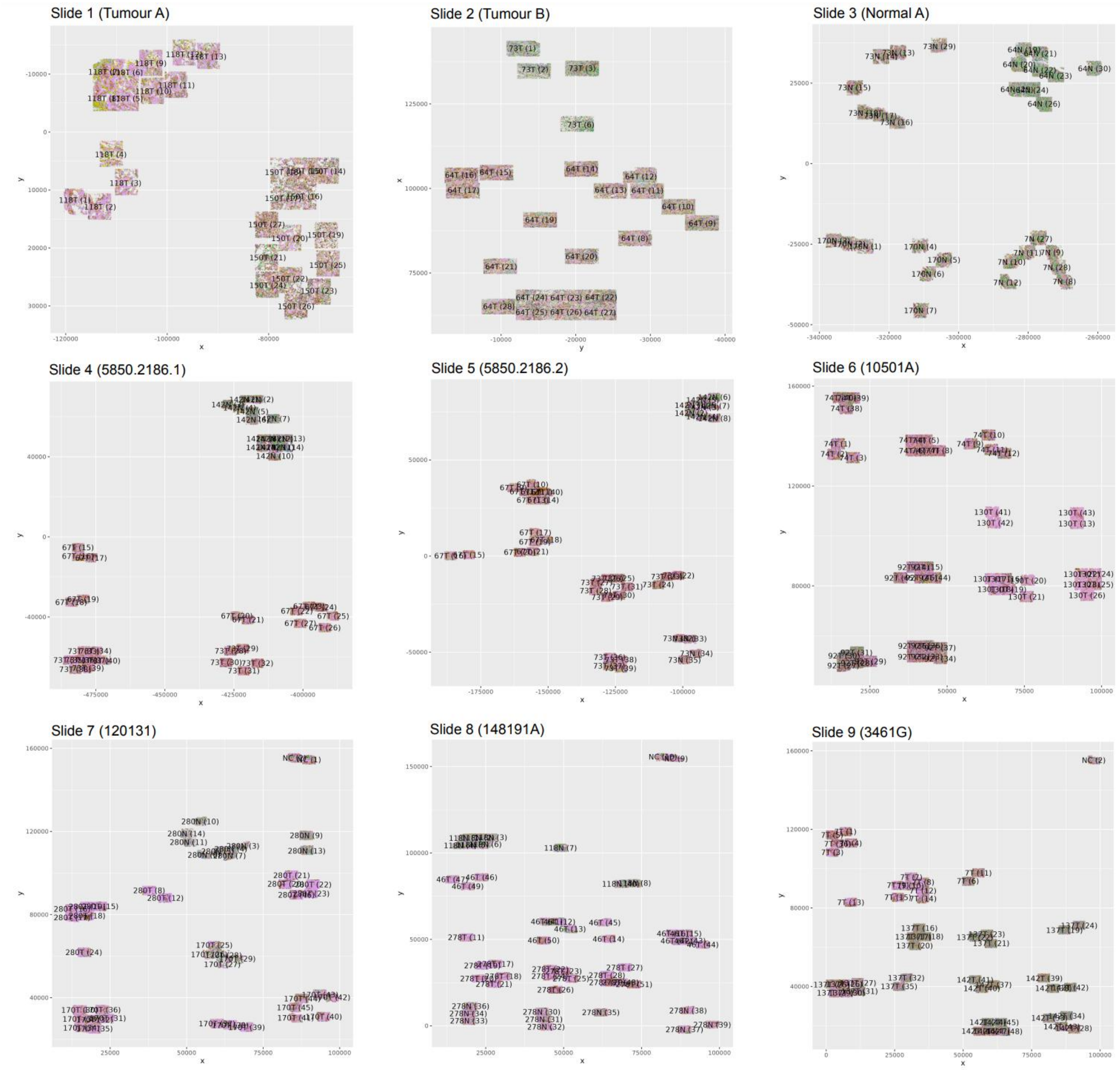


**Figure S1. Layout of the 352 FOVs across the 9 slides of the CosMx run.** FOVs are labelled with sample name and corresponding FOV number.

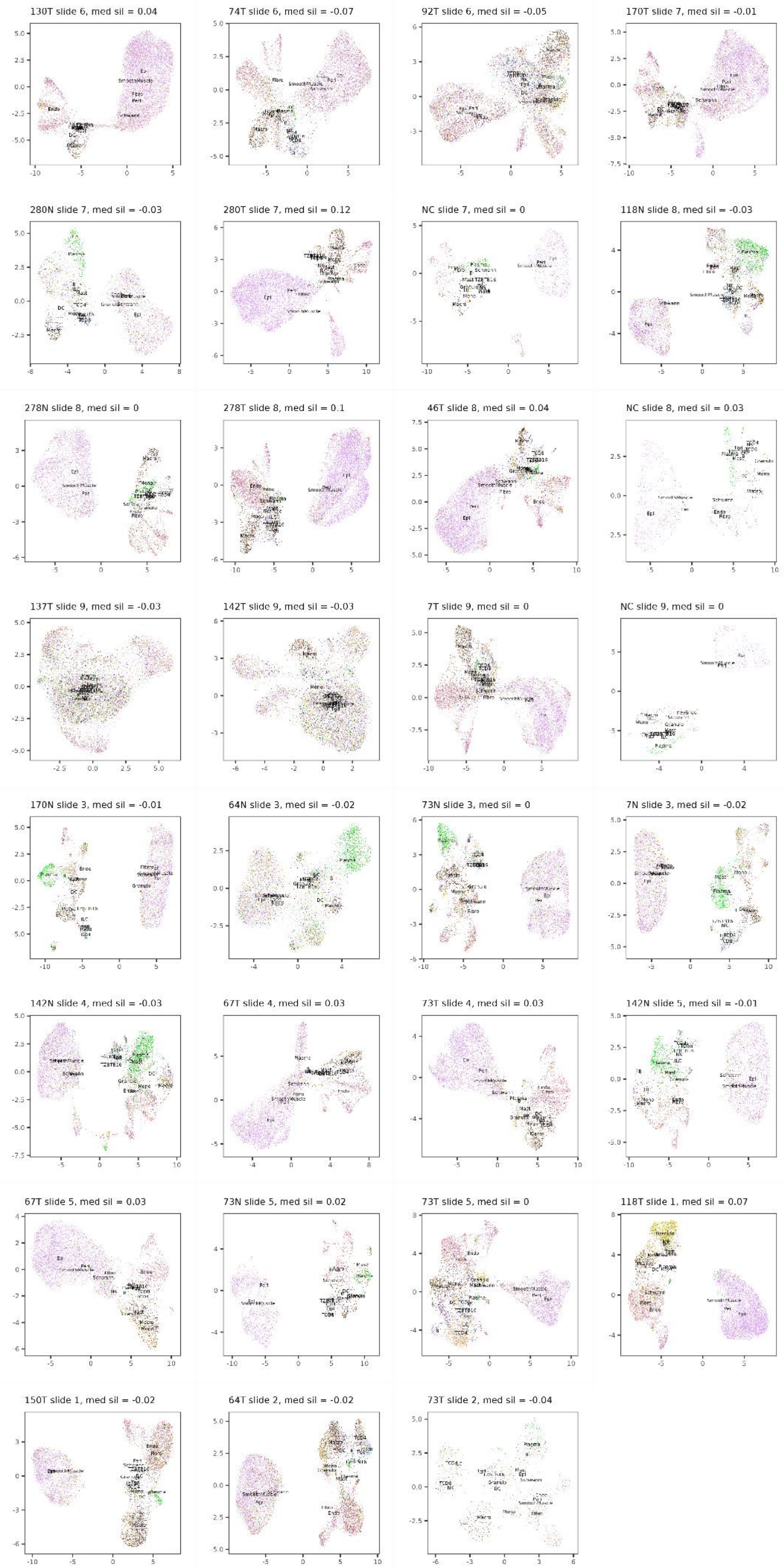
130T slide 6, med sil = 0.04
74T slide 6, med sil = -0.07
92T slide 6, med sil = -0.05
170T slide 7, med sil = -0.01
280N slide 7, med sil = -0.03
280T slide 7, med sil = 0.12
NC slide 7, med sil = 0
118N slide 8, med sil = -0.03
278N slide 8, med sil = 0
278T slide 8, med sil = 0.1
46T slide 8, med sil = 0.04
NC slide 8, med sil = 0.03
137T slide 9, med sil = -0.03
142T slide 9, med sil = -0.03
7T slide 9, med sil = 0
NC slide 9, med sil = 0
170N slide 3, med sil = -0.01
64N slide 3, med sil = -0.02
73N slide 3, med sil = 0
7N slide 3, med sil = -0.02
142N slide 4, med sil = -0.03
67T slide 4, med sil = 0.03
73T slide 4, med sil = 0.03
142N slide 5, med sil = -0.01
67T slide 5, med sil = 0.03
73N slide 5, med sil = 0.02
73T slide 5, med sil = 0
118T slide 1, med sil = 0.07
150T slide 1, med sil = -0.02
64T slide 2, med sil = -0.02
73T slide 2, med sil = -0.04

**Figure S2. Single sample clustering, annotation and QC.** Each sample from each slide was QC filtered, clustered using Seurat (V5) and annotated using singleR. Median silhouette scores were calculated from the cell type annotations compared against the PCA embeddings, similar to the method employed by Stuart *et al*.[27]. The silhouette score is between -1 and 1 and calculates how similar a given cell is to cells in its own cluster vs other clusters.

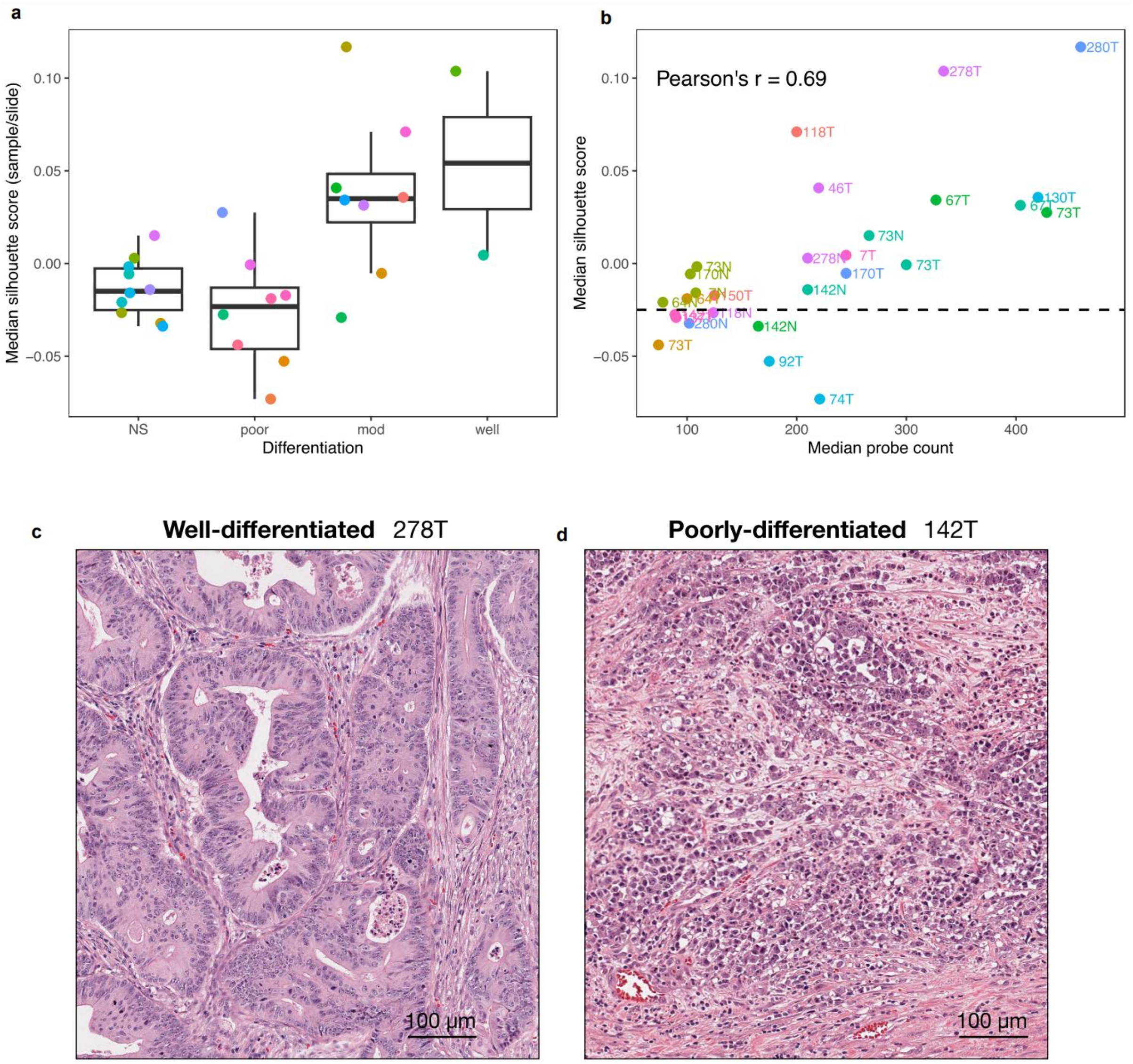


**Figure S3. Silhouette scores reflect read counts and tissue differentiation. a.** Silhouette scores vs pathologist identified tumour differentiation status. NS are normal samples. **b.** Median probe counts correlated with median silhouette score (Pearson's r = 0.69). Based on visual examination of clustering of CosMx slides, samples with a median silhouette score < -0.025 were considered low quality (dotted line). **c**. Representative H&E stain of a well-differentiated tumour. **d**. Representative H&E stain of a poorly-differentiated tumour.

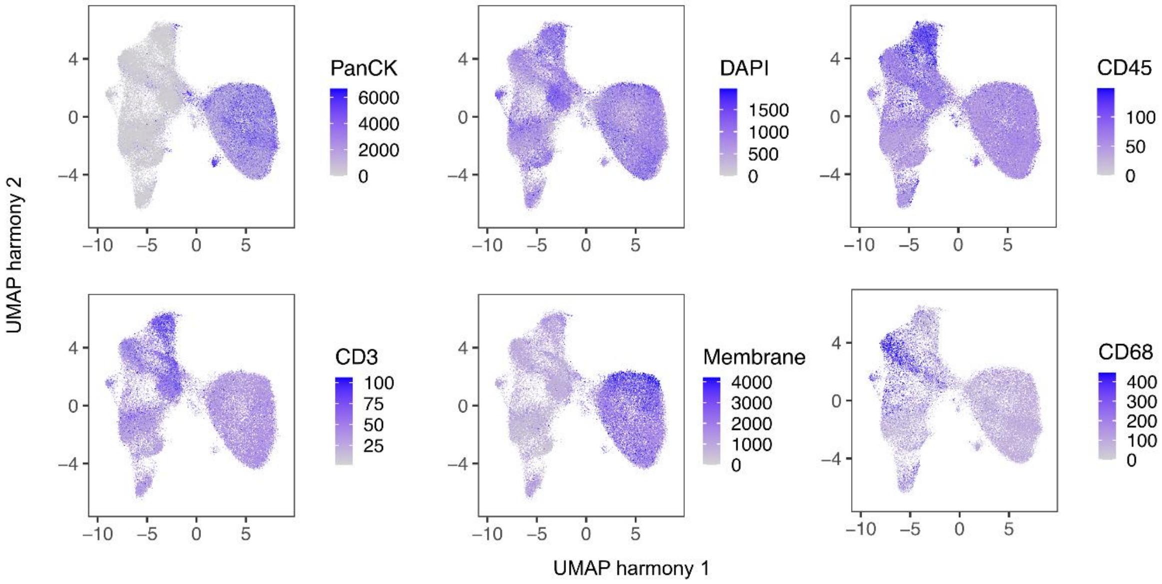


**Figure S4. IF protein detection aligns with annotated cell type (V1 probeset).**

Mean marker expression for PanCK (epithelial), DAPI (nuclear), CD45 (immune), CD68 (myeloid), CD3 (T cells) and membrane stain.

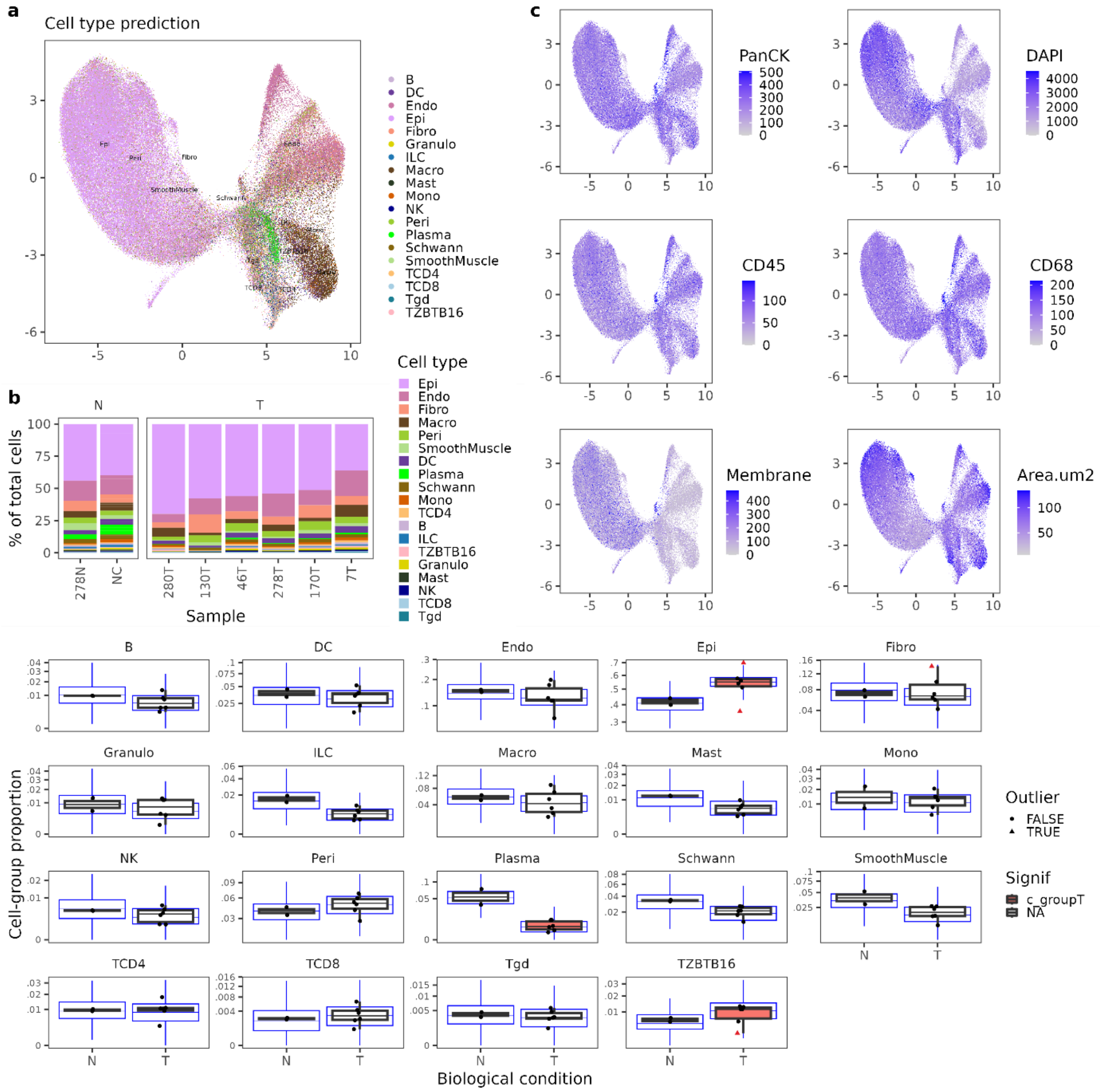


**Figure S5. Annotation and composition of high quality cells (V2 probeset). a.** UMAP of harmony integrated high quality cells. **b.** The cellular composition of each sample split by tumour and normal. **c.** Mean marker expression for PanCK (epithelial)**,** DAPI (nuclear), CD45 (immune), CD68 (myeloid), membrane stain and total cellular area. **d.** Changes in cell type proportions (DA) between tumour and normal samples as calculated by sccomp, colour indicates a significant association of sample type with cell composition, the blue boxplots represent a posterior predictive check.

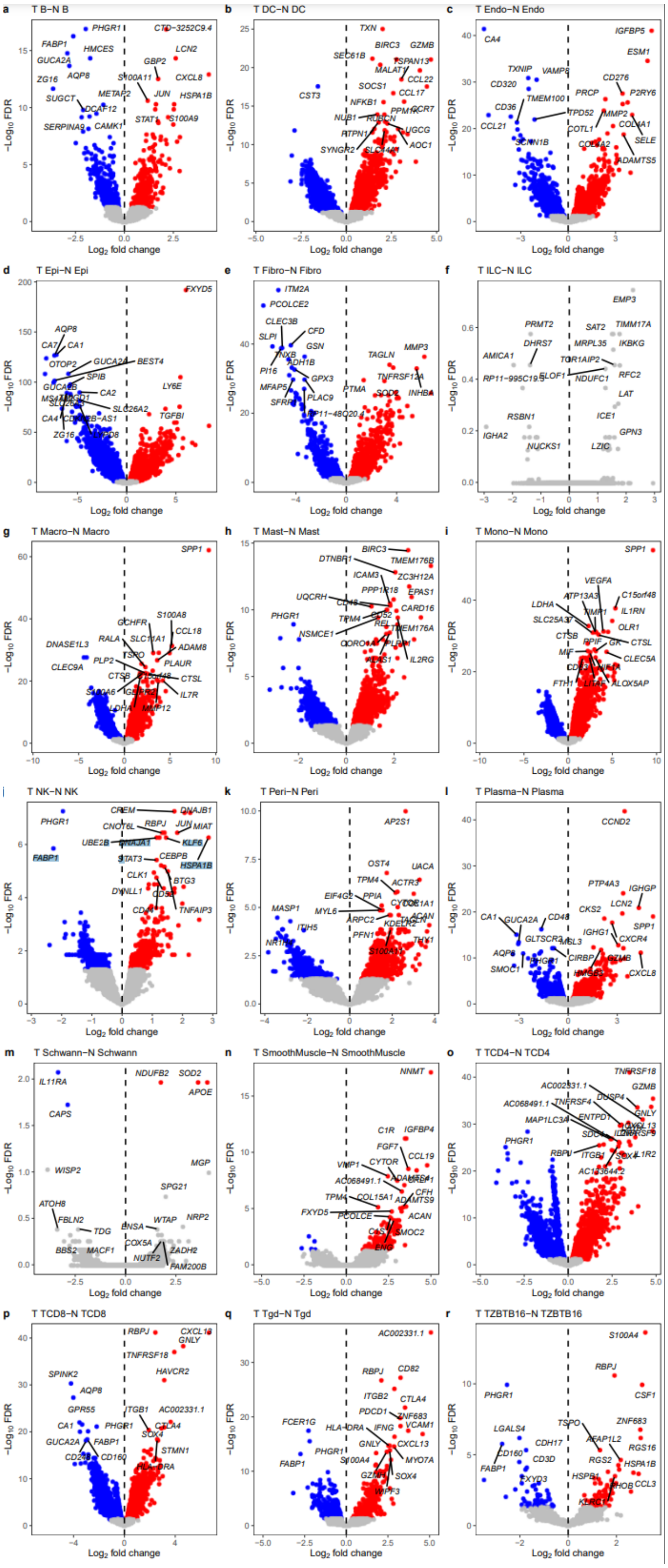


**Figure S6. MMR atlas tumour vs normal. a-r**. Volcano plots of comparing tumour and normal for each cell type in the MMR atlas.

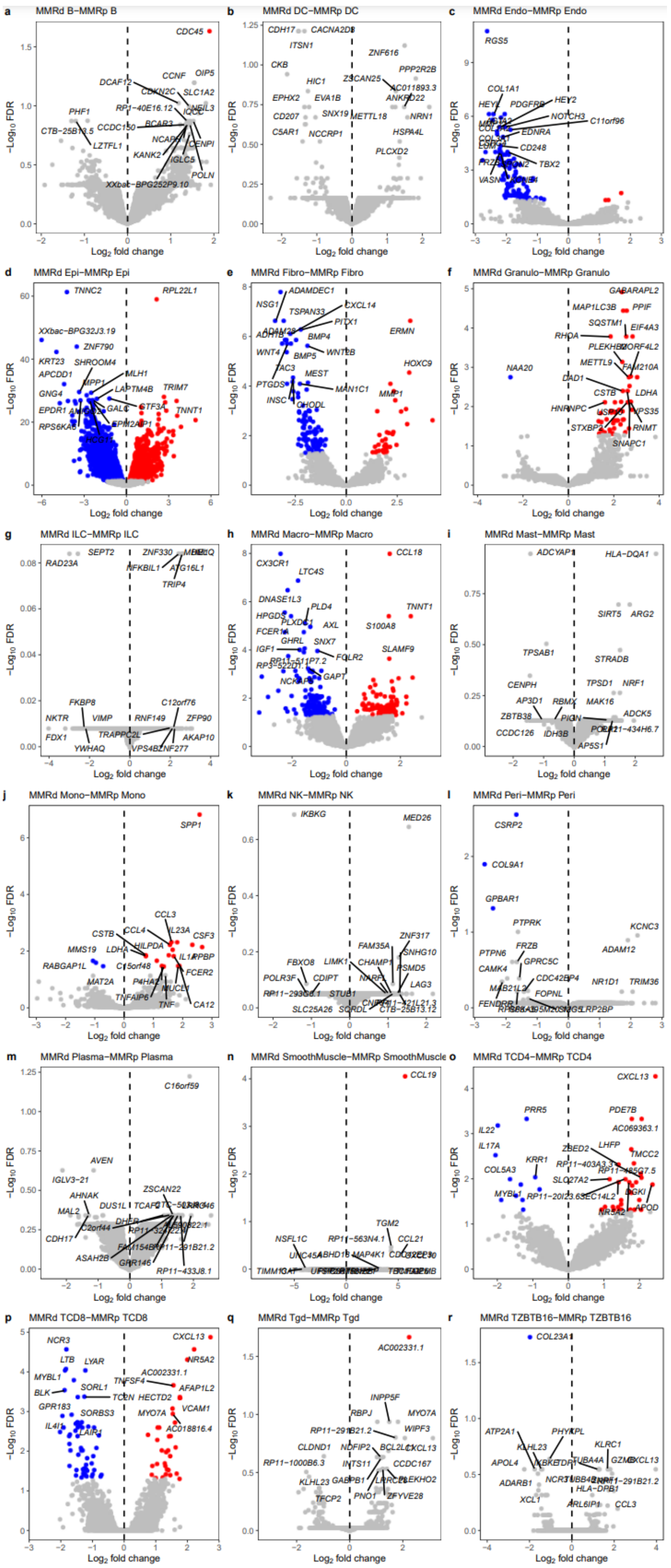


**Figure S7. MMR atlas MMRd tumours vs MMRp tumours. a-r**. Volcano plots comparing MMRd tumours to MMRp tumours for each cell type in the MMR atlas.

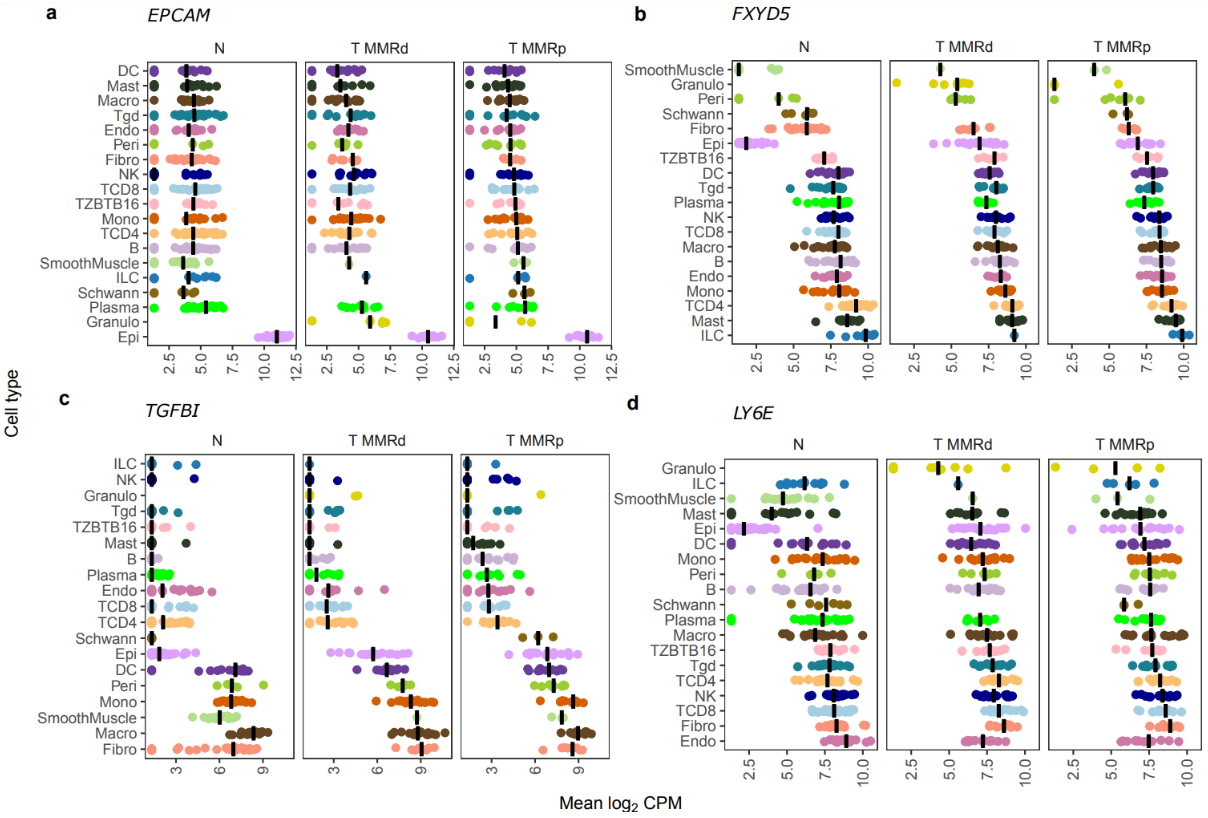


**Figure S8. Key CRC-associated gene expression in all cell types. a-f**. Gene expression (pseudobulked $log_2$ CPM) of key genes split into tumour MMRp/d and normal groupings. The line represents the median expression of each cell type.

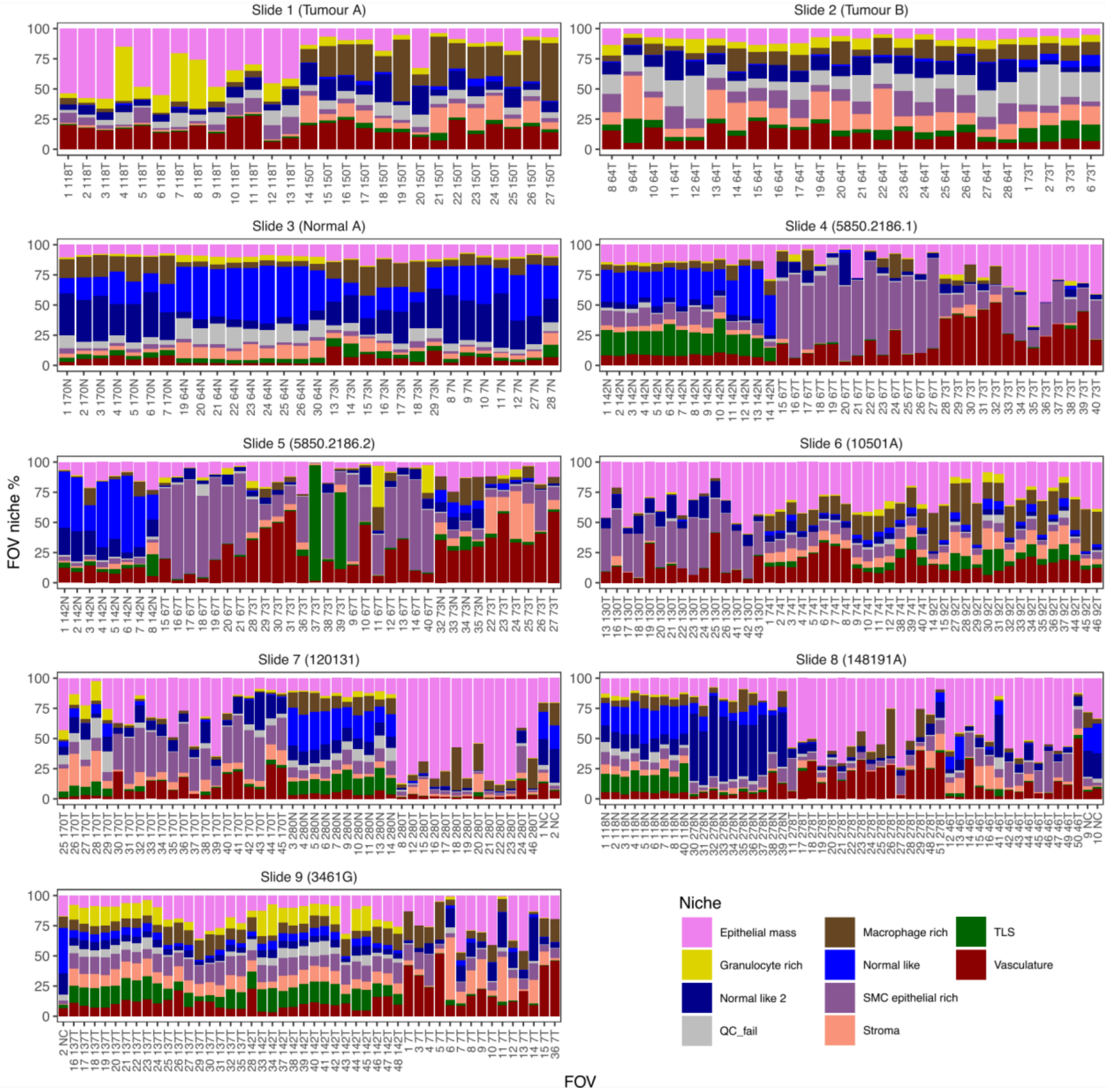


**Figure S9. Per-FOV niche composition.** The cellular nice proportions of each FOV analysed by CosMx, split by slide and ordered by sample.

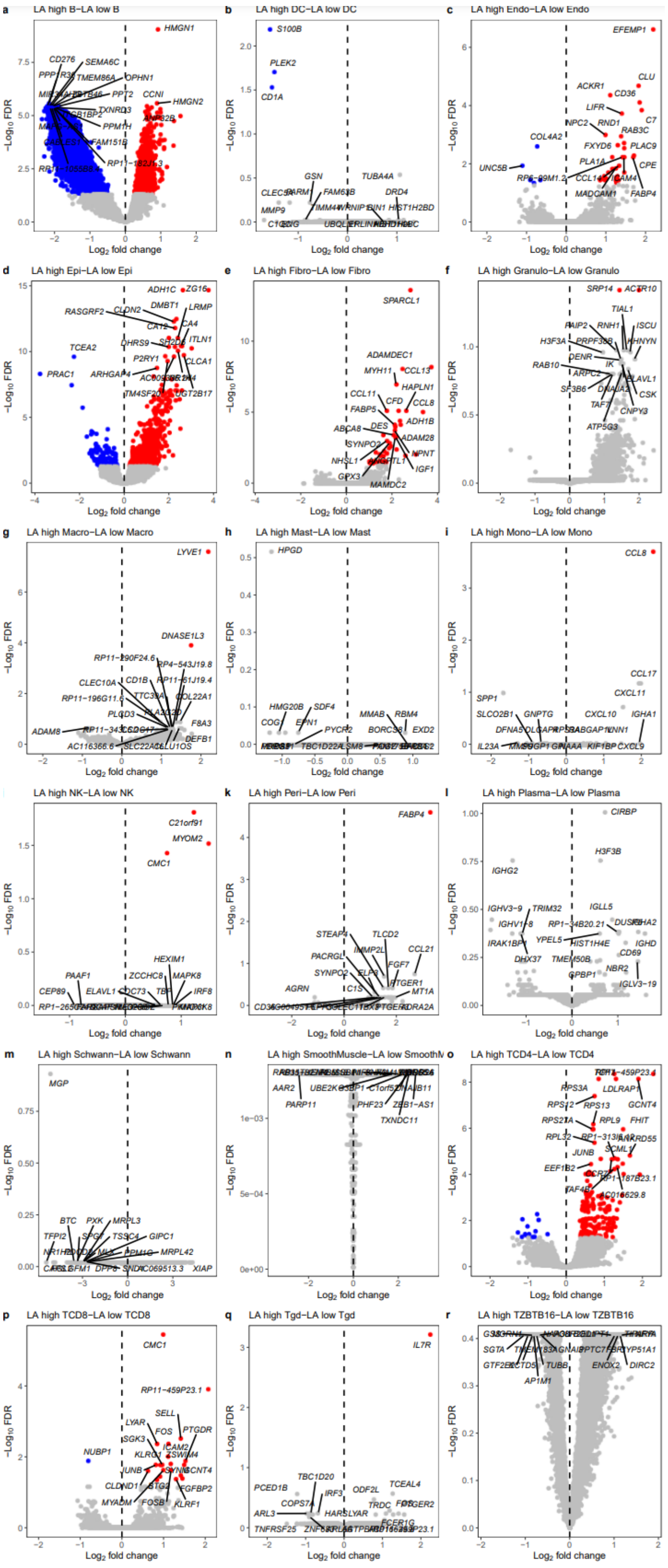


**Figure S10. MMR atlas LA high vs LA low tumours. a-r**. Volcano plots comparing tumours with high *SELL*/*TCF7* CD4 T cell infiltration (LA high) tumours vs low for each cell type in the MMR atlas.

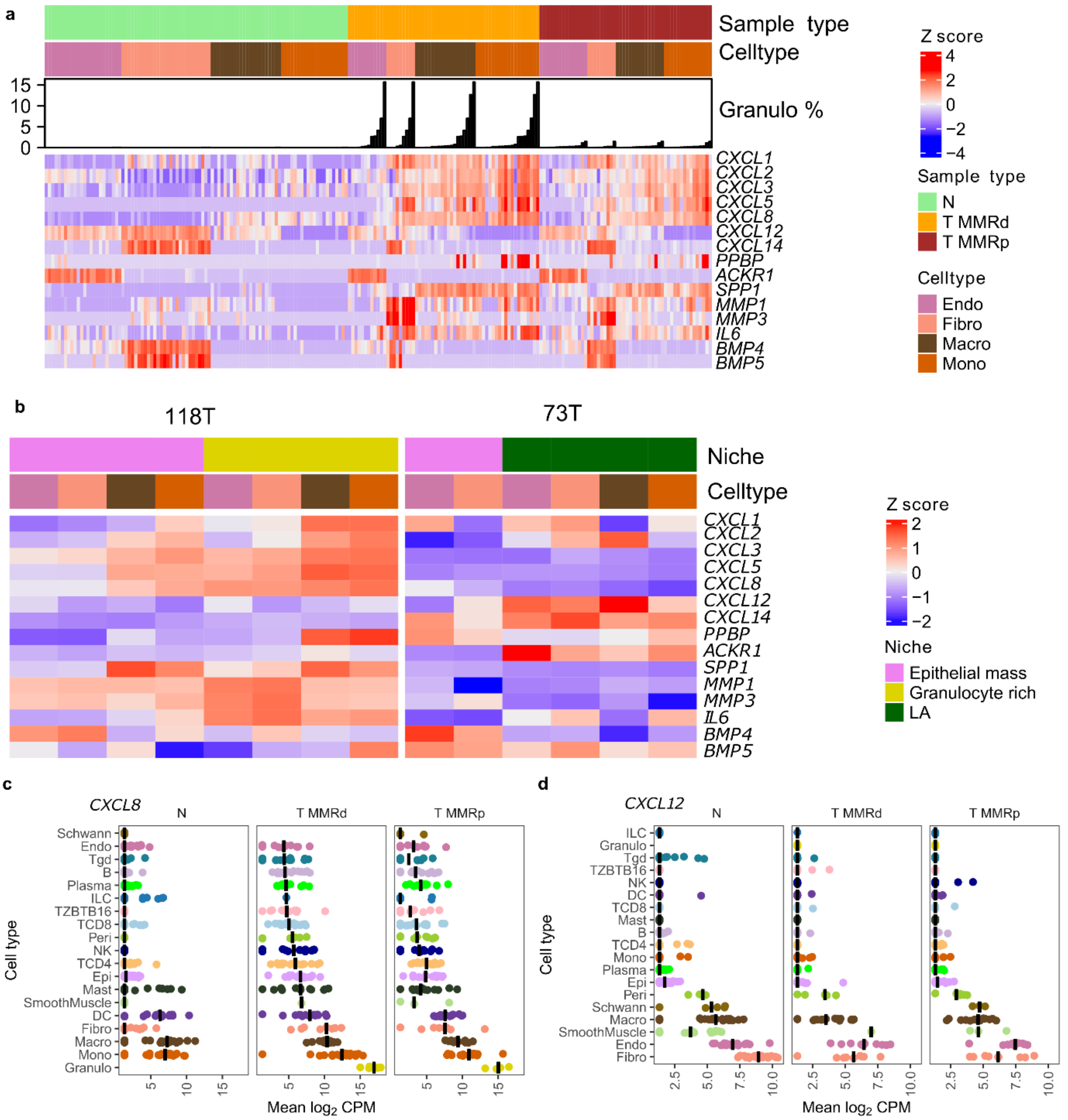


**Figure S11. Key neutrophil chemoattractants. a.** Key fibroblast/myeloid gene expression in the MMR atlas. **b**. The expression of the same key genes as a in the epithelial and granulocyte niches of sample 118T and the epithelial and LA niches of sample 73T**. c** Gene expression (pseudobulked $log_2$ CPM) of *CXCL8* and *CXCL12* (**d**) split into tumour MMRp/d and normal groupings.

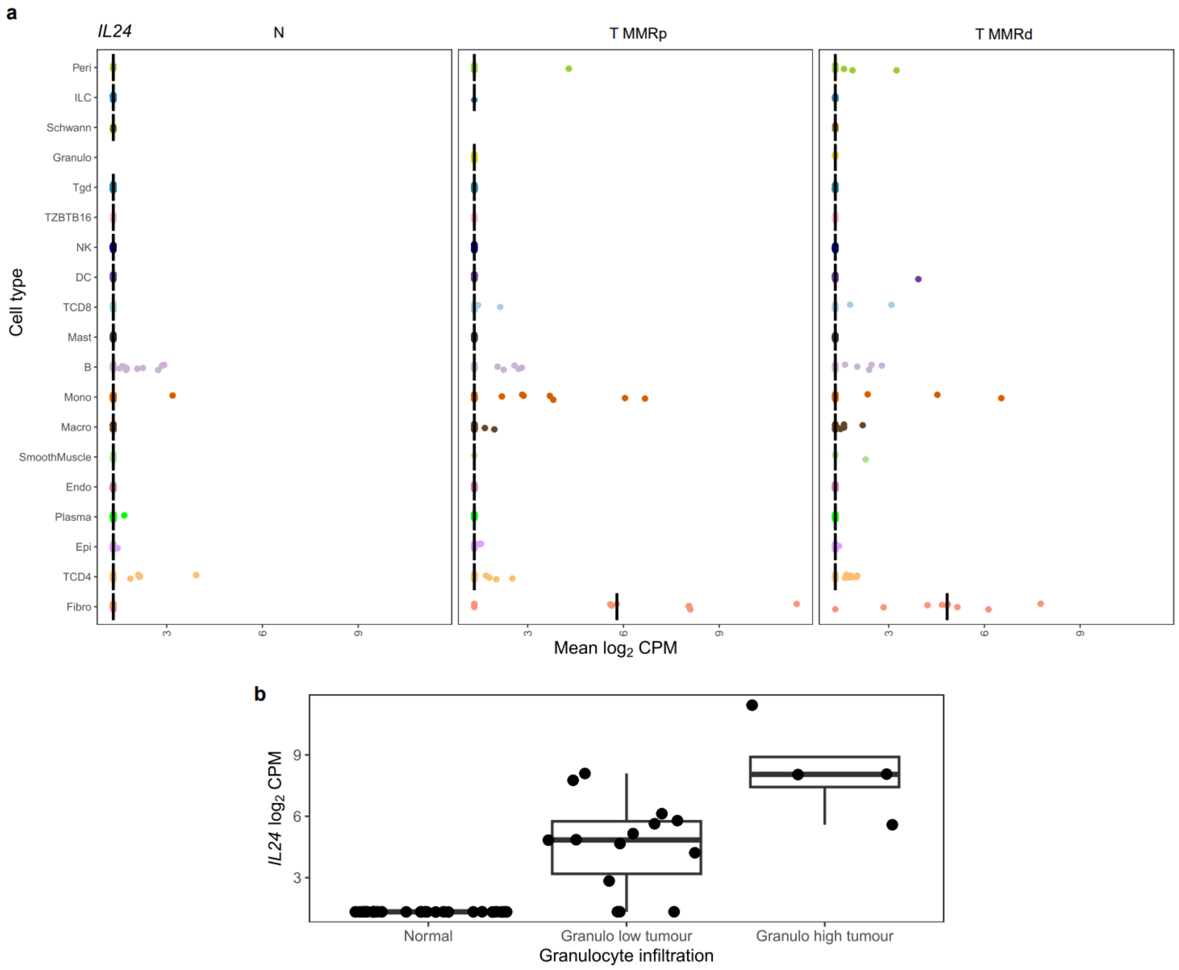


**Figure S12. IL24 expression in normal, tumour and neutrophil-high MMR atlas samples. a** Expression of *IL24* from normal colon and MMRp/d tumours.

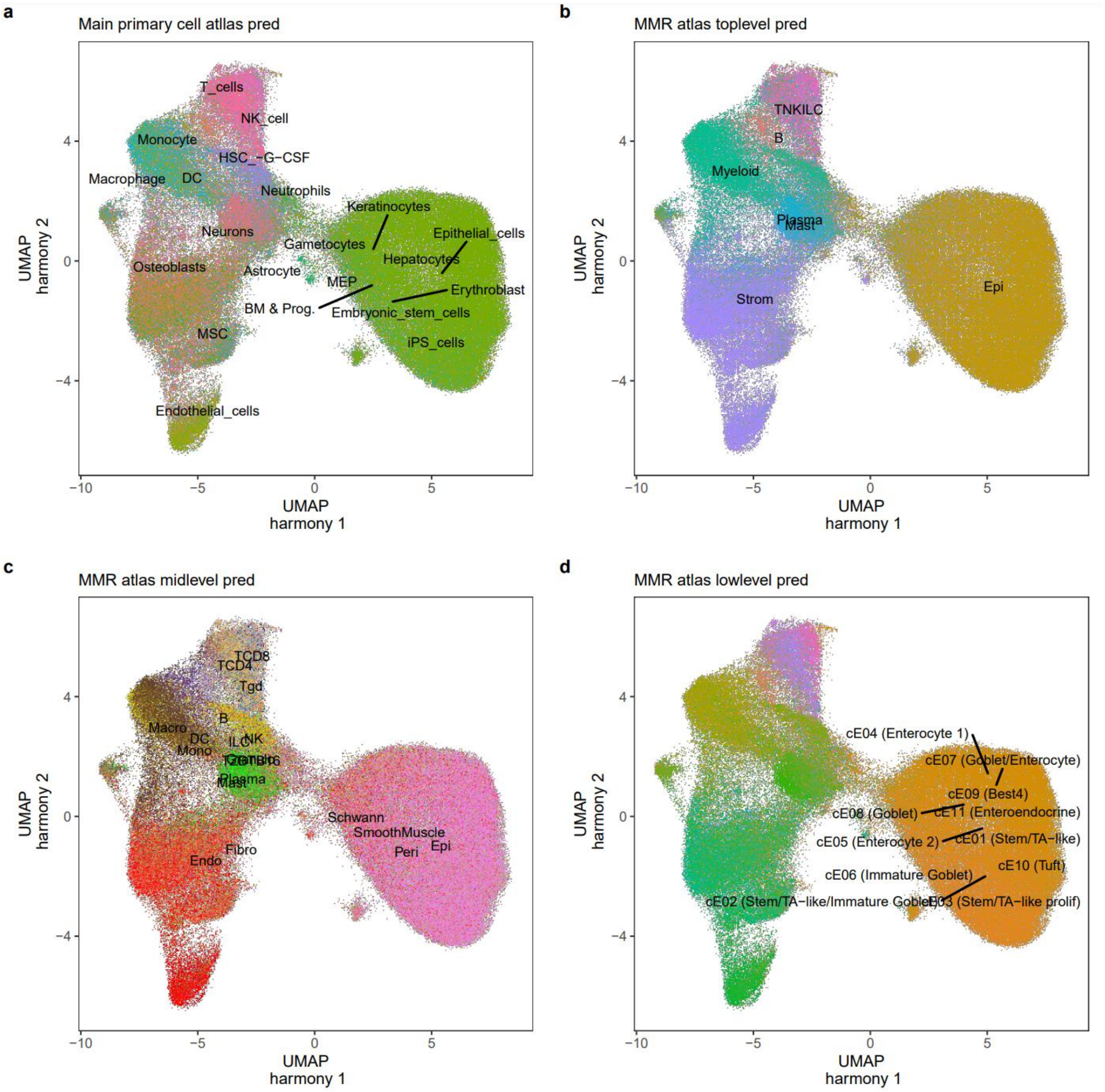


**Figure S13. SingleR cell type predictions (V1 probeset).** Predictions were generated using the human primary cell atlas (Mabbott *et al.*[74]) main labels (**a**) or top-level (**b**), mid-level (**c**) and low-level (**d**) MMR atlas predictions.

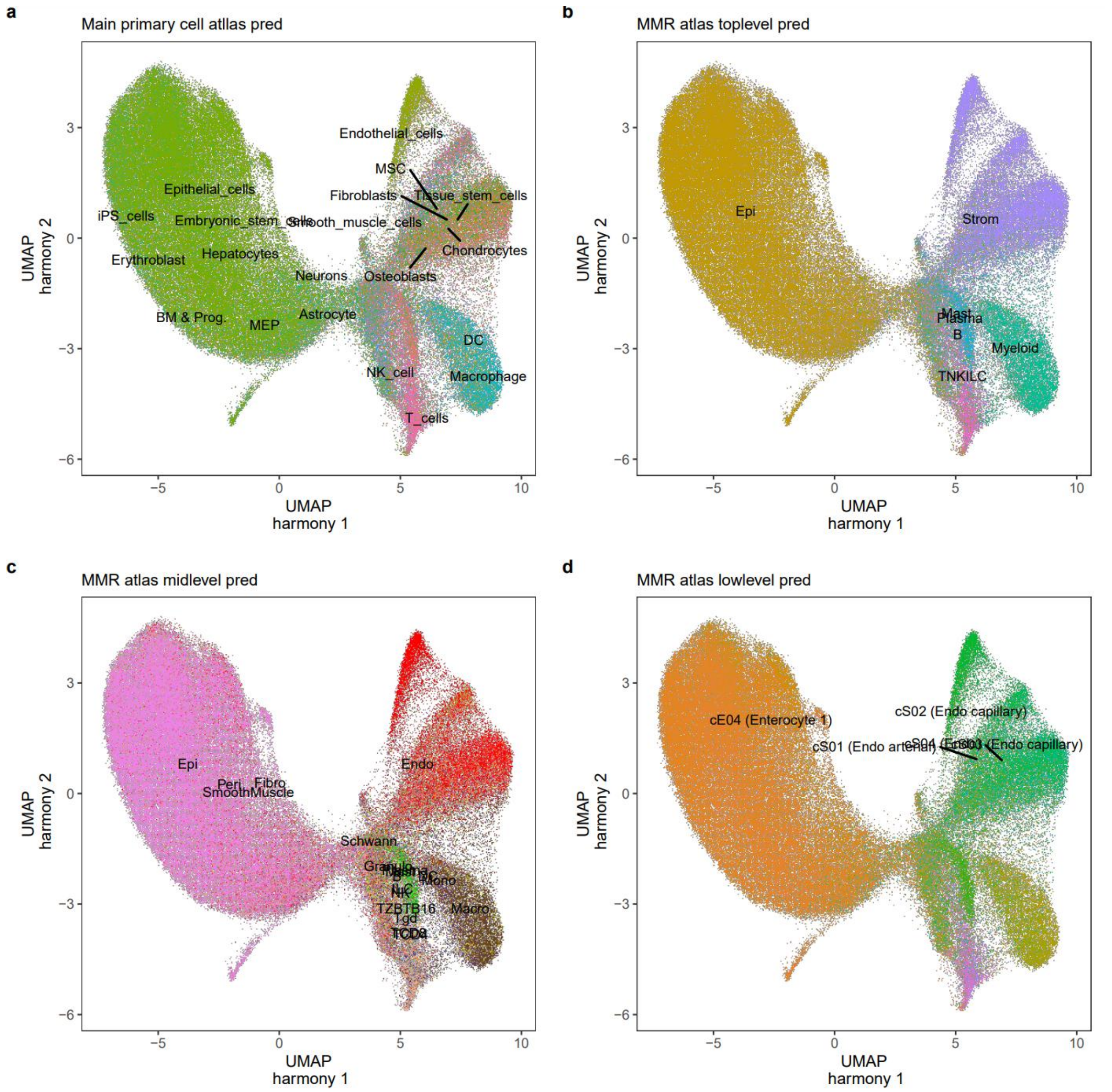


**Figure S14. SingleR cell type predictions (V2 probeset).** Predictions were generated using the human primary cell atlas (Mabbott *et al.*[74]) main labels (**a**) or top-level (**b**), mid-level (**c**) and low-level (**d**) MMR atlas predictions.

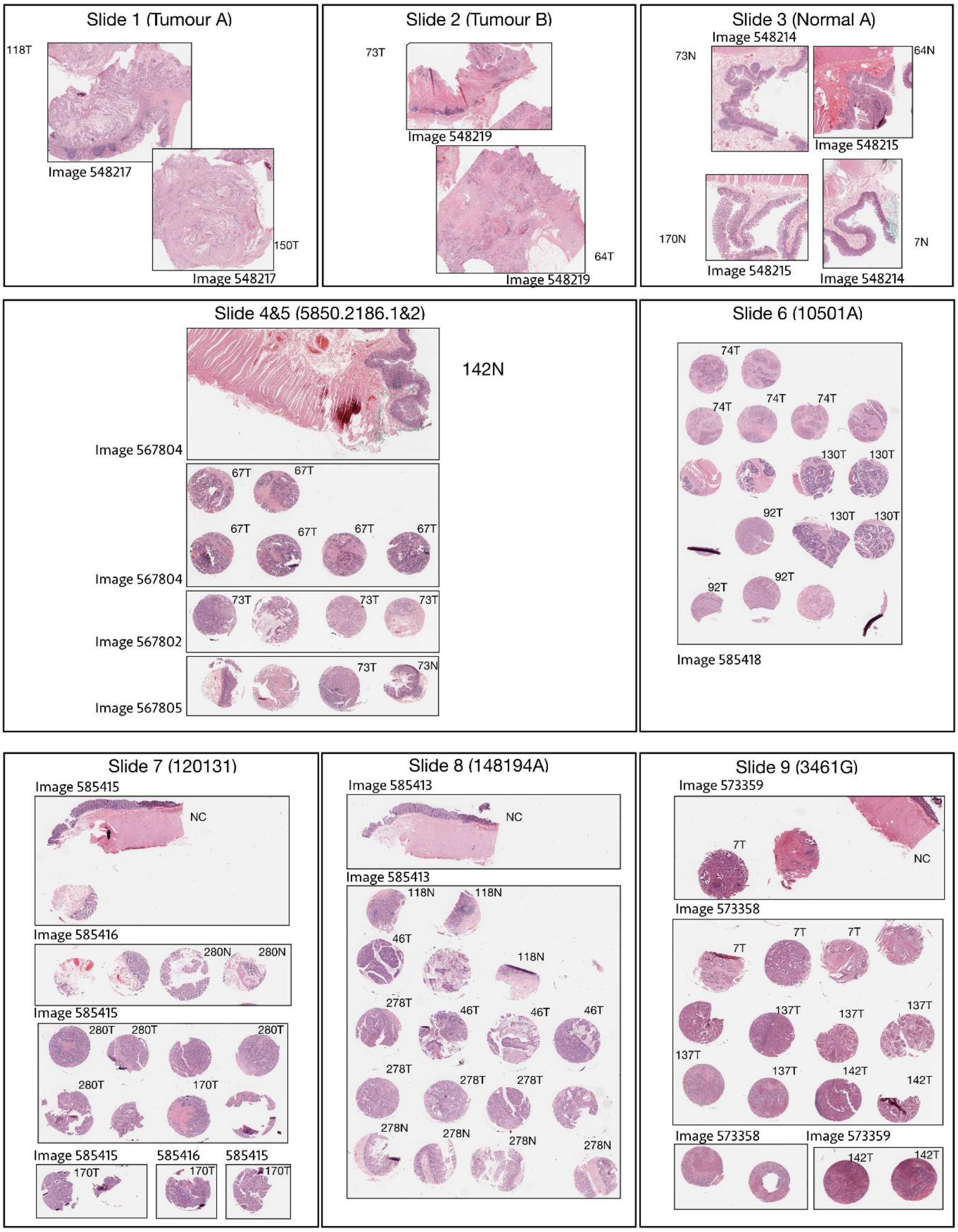


**Figure S15. High resolution H & E images of CosMx slides.** H & E images of the slide layouts that were used for the CoxMx experiments.

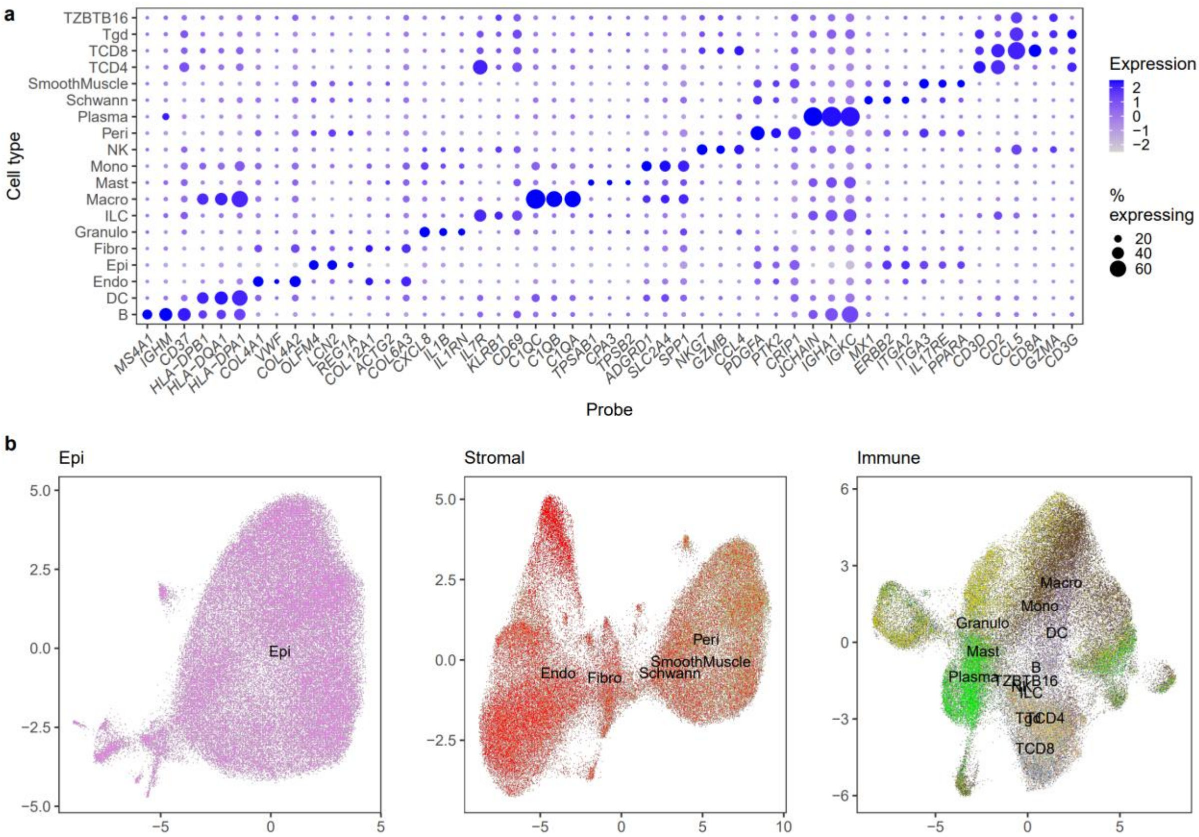

**Figure S16. SingleR annotated cell markers and subclustering (V1 probeset). a** Dotplot of the top 3 marker genes (by $\log_2$ fold change) of each cell type. **b.** Subclusterining of the epithelial, immune and stromal components.

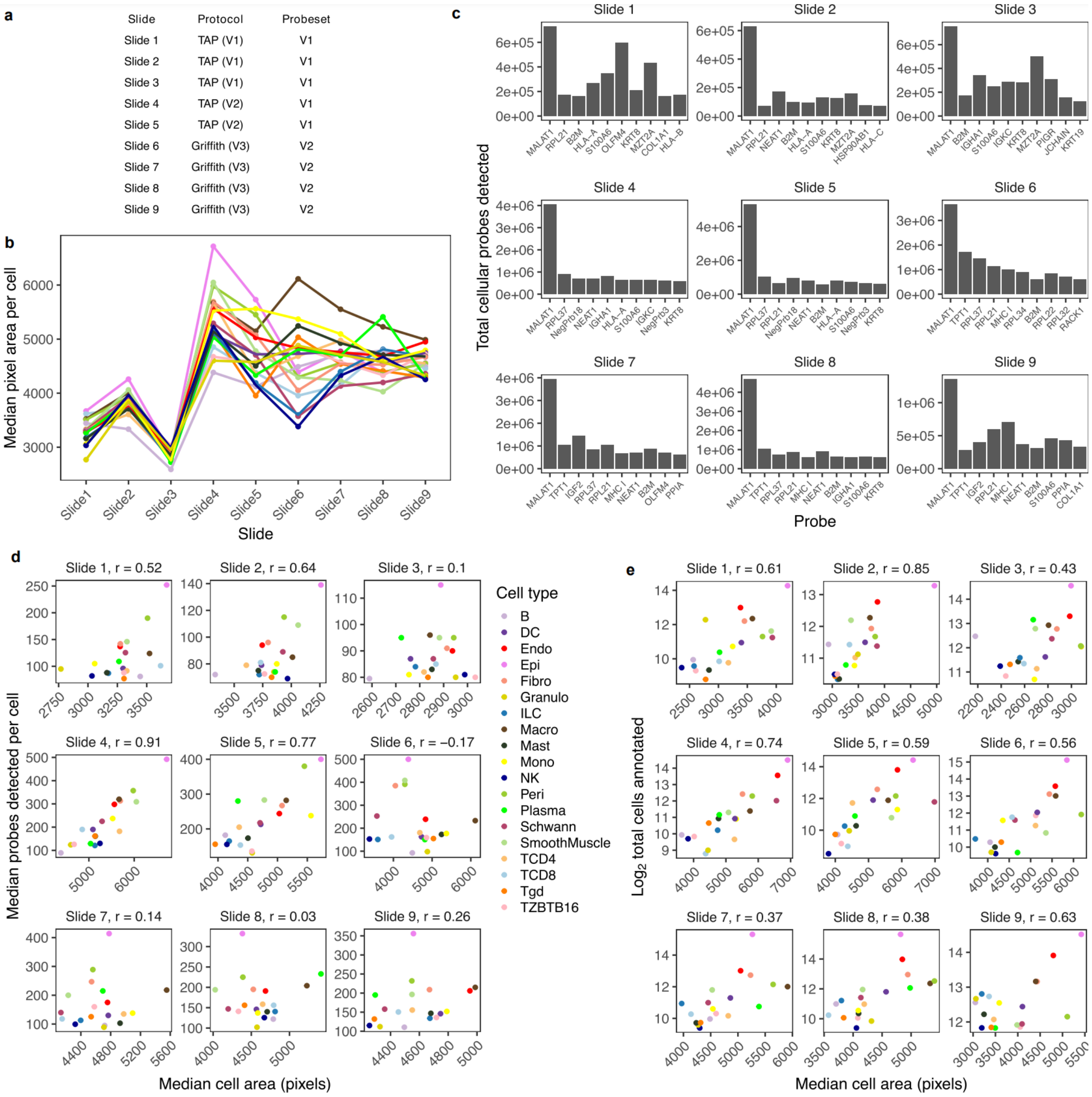


**Figure S17. The effect of probe and cell size on cell type annotation across CosMx version. a.** The CosMx versions used to run each slide. **b.** The median pixel are per cell for each slide, split by annotated cell type. **c.** Median cell area vs median probes detected in each cell type (grouped by slide). Cell size is highly correlated with the number of cells detected on slides 1, 2, 4 and 5. **d.** Top 10 most commonly detected probes on each slide. **e.** Cell area vs the number of cells detected. In general, the larger a cell is, the more cells of that type are detected.

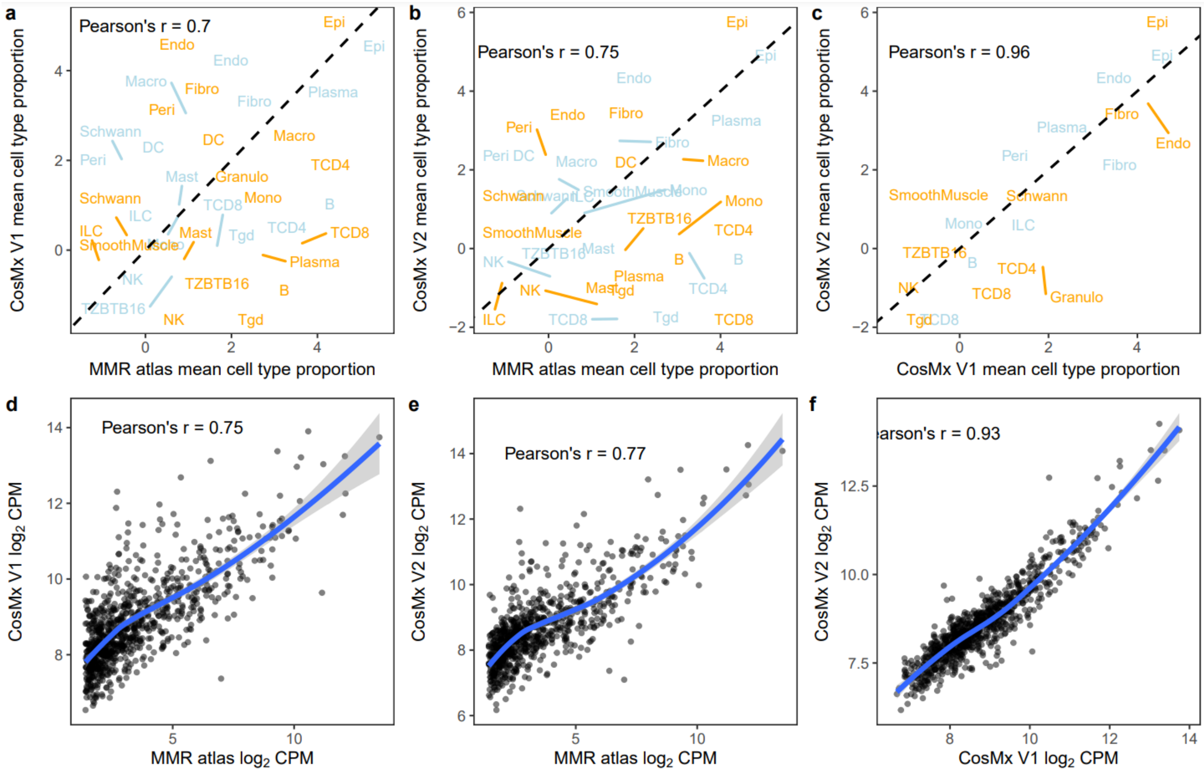

**Figure S18. Comparing CosMx counts and cellular abundance with scRNA-Seq.** Correlation of the average cell type proportions for tumour (orange) and normal (light blue) between the MMR atlas and the V1 CosMx probeset (**a**) and the V2 CosMx probeset (**b**) and V1 vs V2 (**c**). **d-f.** The same contrasts as **a-c** comparing gene detection of commonly expressed genes/detected probes.

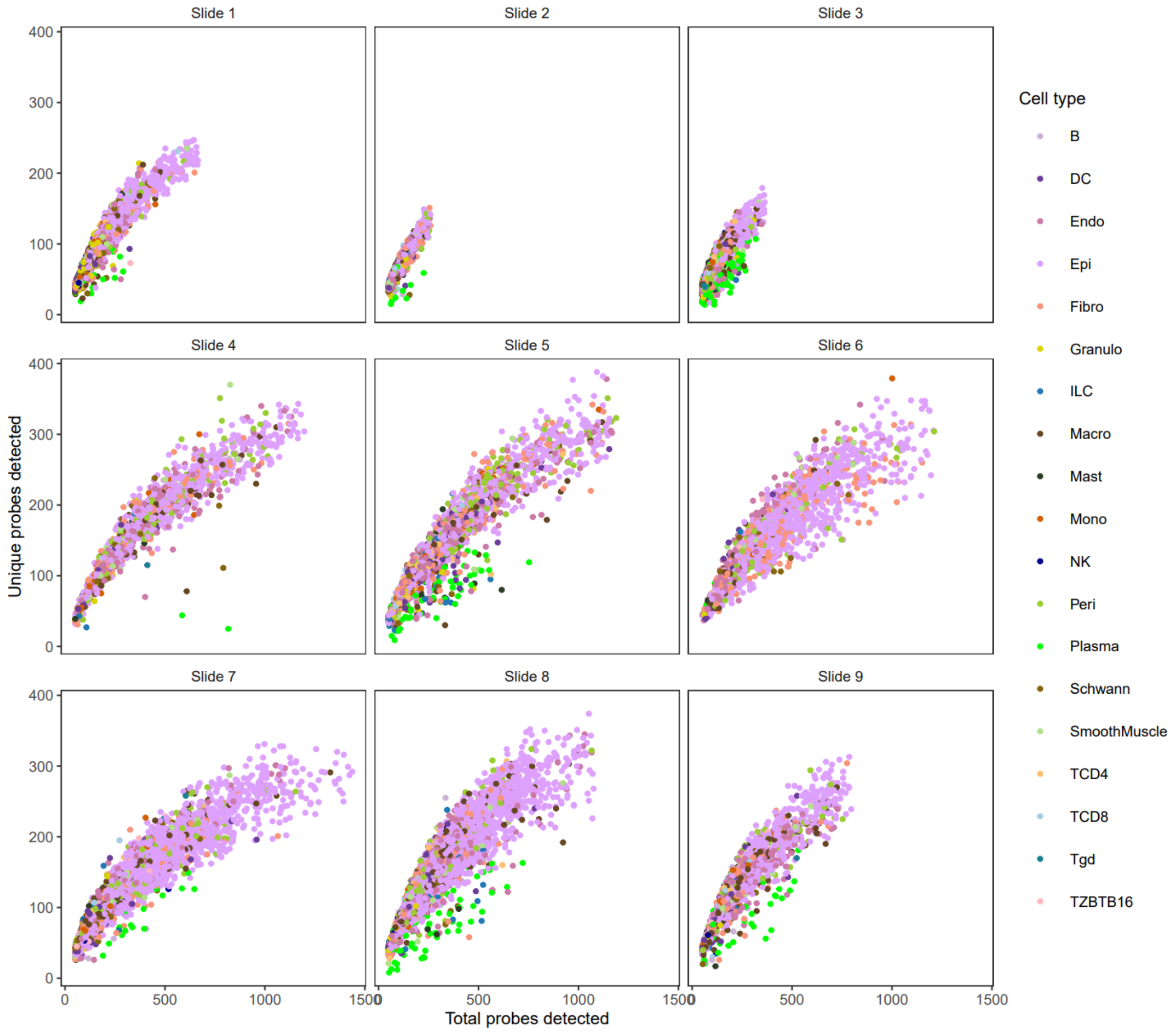


**Figure S19. Scatterplot of the number of probes detected vs the unique probes detected.** 10,000 cells were sampled from each slide and plotted. More detected probes consistently results in more unique genes detected suggesting saturation has not been achieved.

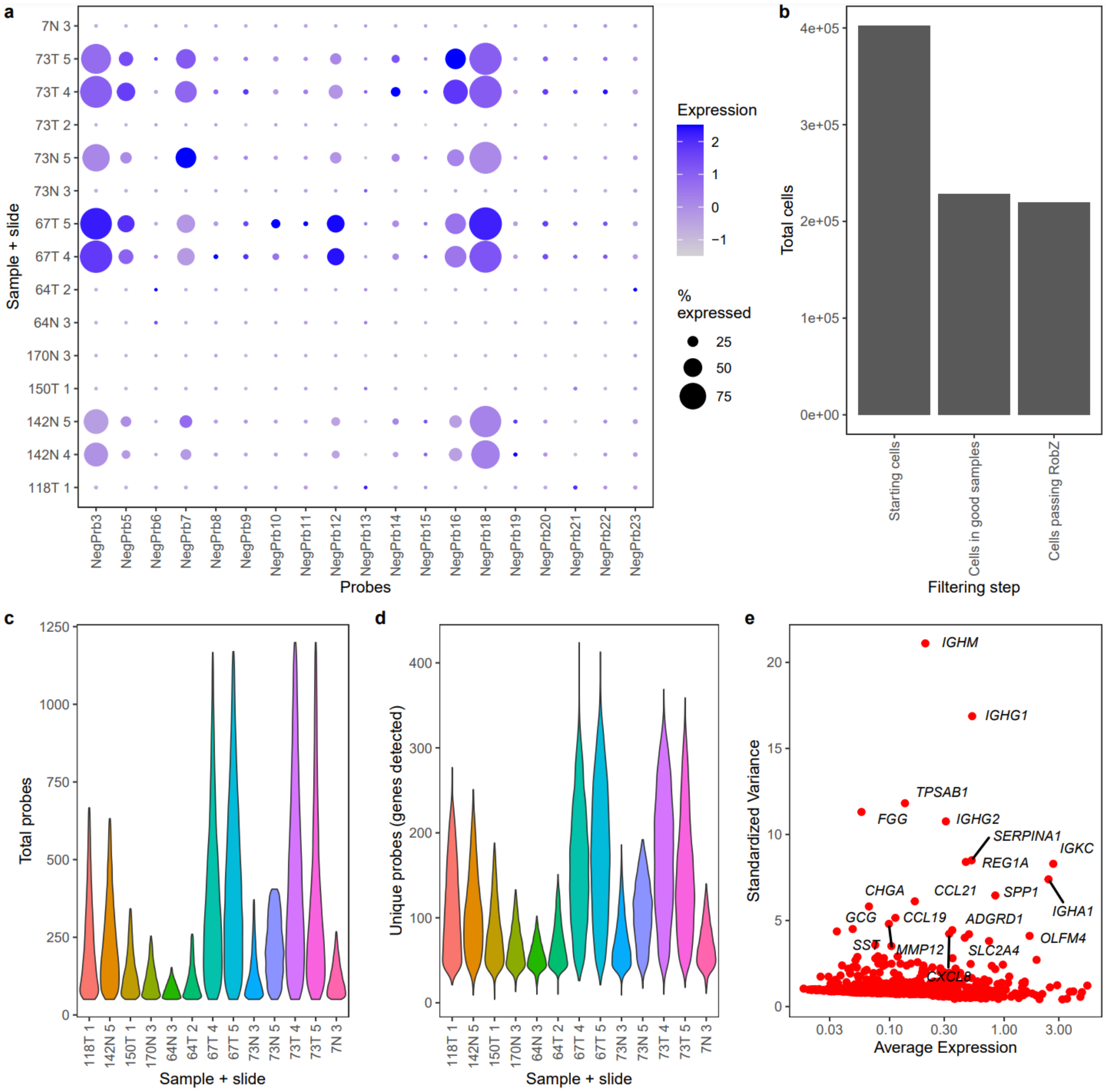


**Figure S20. CosMx V1 QC and filtering. a** Dot plot of negative probe expression in cells from each sample/slide. Probes 3, 5, 7, 12, 16 and 18 appear to not be truly negative on slides 4 and 5. **b.** Cells remaining at each stage of QC filtering. **c.** Violin plots of median probe counts per cell and unique probes detected in each cell (**d**) following QC filtering. **e.** Top variable genes vs average expression in post-QC cells.

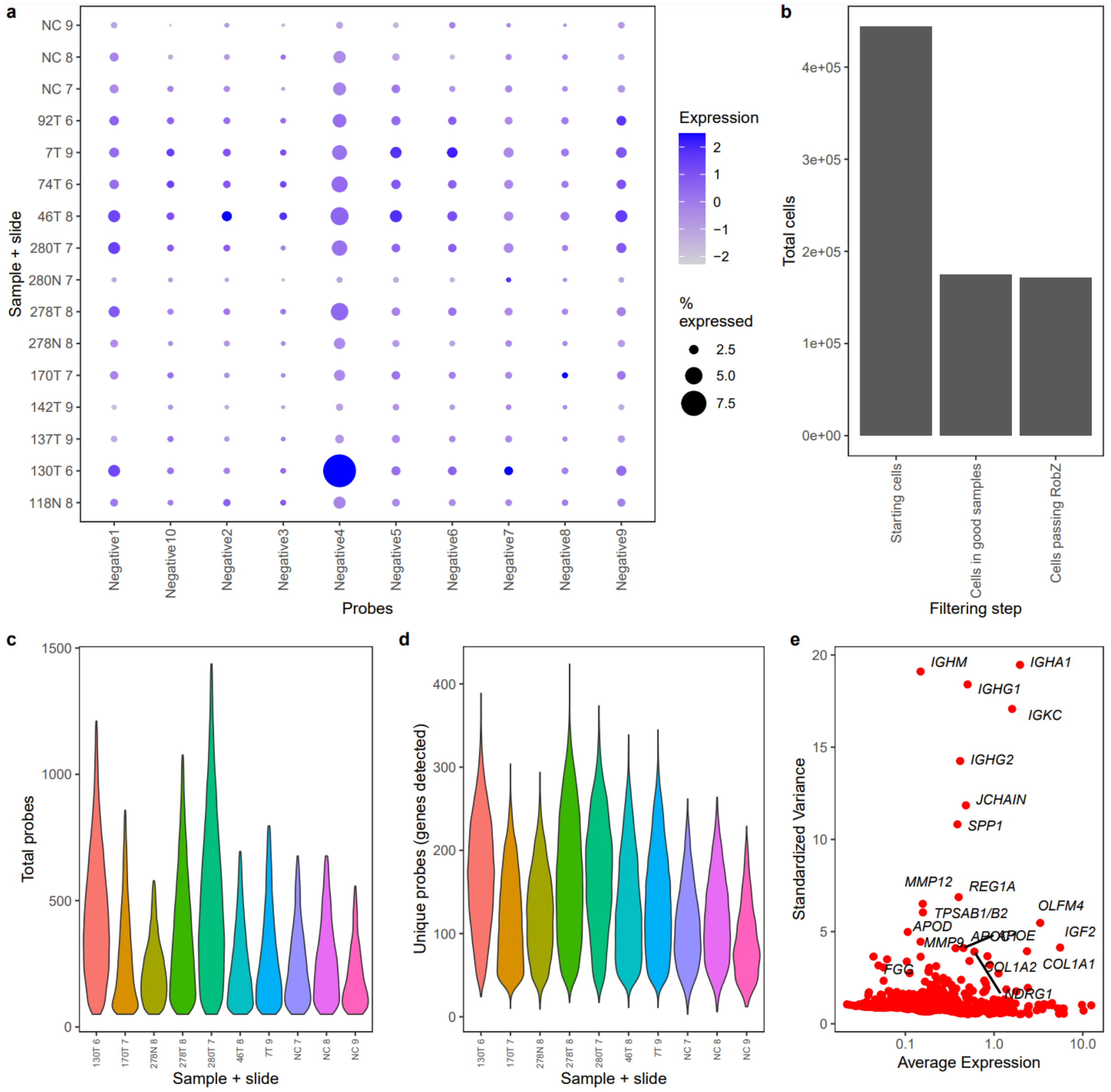


**Figure S21. CosMx V2 QC and filtering. a** Dot plot of negative probe expression in cells from each sample/slide. **b.** Cells remaining at each stage of QC filtering. **c.** Violin plots of median probe counts per cell and unique probes detected in each cell (**d**) following QC filtering. **e.** Top variable genes vs average expression in post-QC cells.

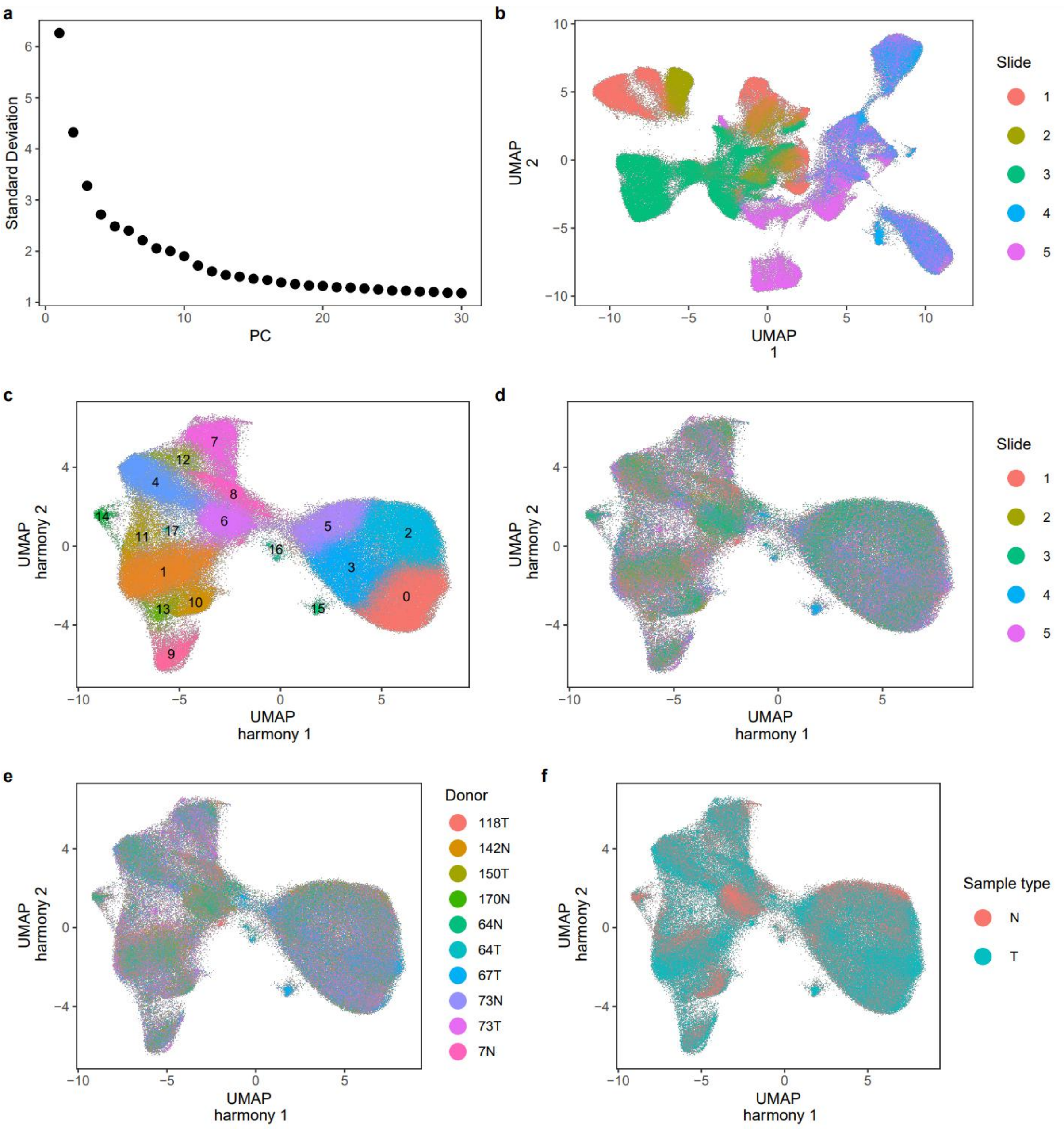


**Figure S22. CosMx V1 clustering and integration. a.** Elbow plot based on variance explained by each of the first 30 PCs calculated from the high quality V1 samples. **b.** UMAP of cell types coloured by cluster. A clear slide level batch effect can be observed. **c.** UMAP of harmony integrated cells removing sample and slide level batch effects coloured by seurat cluster. **d-f.** UMAP of integrated cells coloured by slide (**d**), sample (**e**) and sample type (**f**).

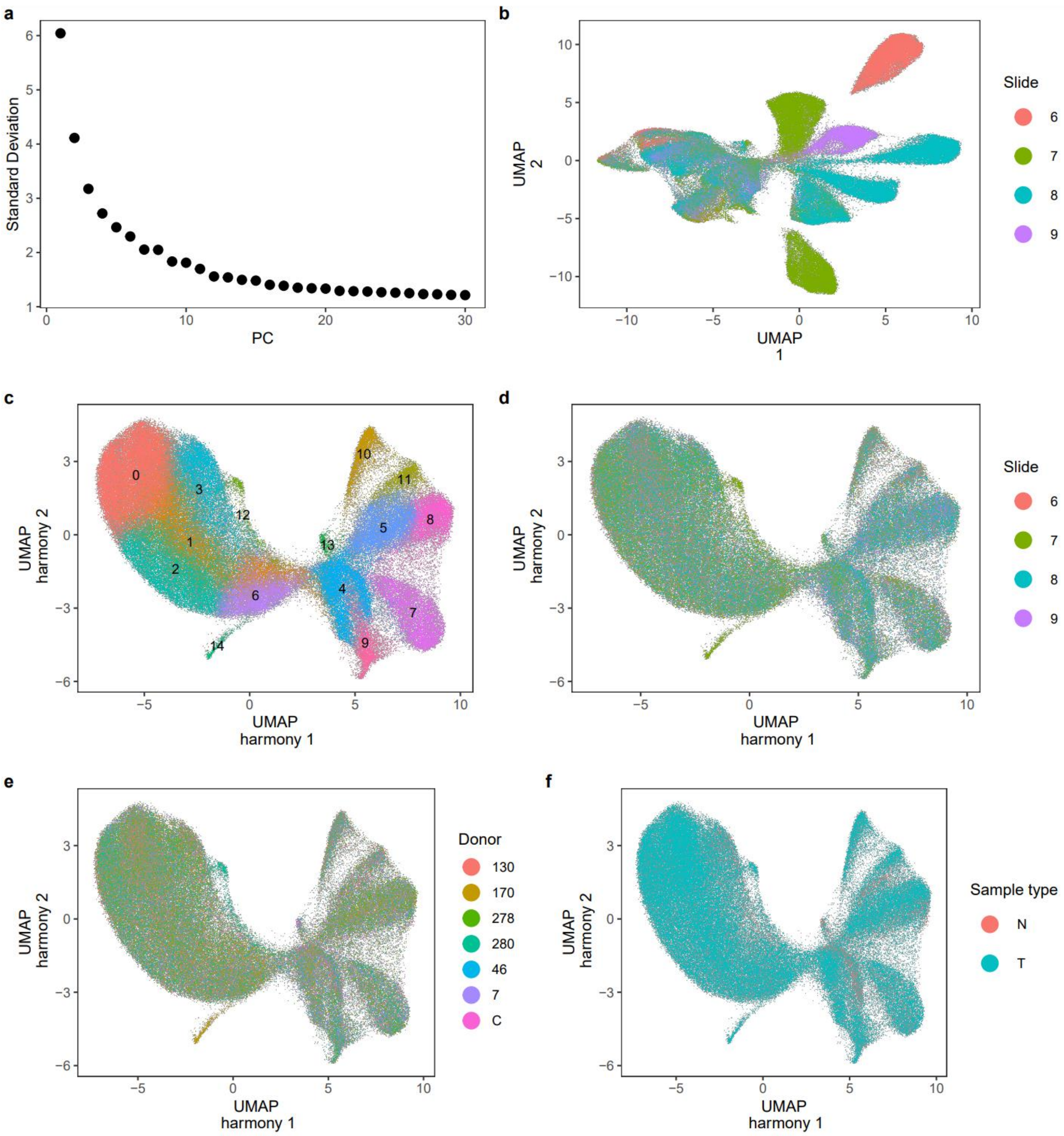


**Figure S23. CosMx V2 clustering and integration. a.** Elbow plot based on variance explained by each of the first 30 PCs calculated from the high quality V2 samples. **b.** UMAP of cell types coloured by cluster. **c.** UMAP of harmony integrated cells removing sample and slide level batch effects coloured by seurat cluster. **d-f.** UMAP of integrated cells coloured by slide (**d**), sample (**e**) and sample type (**f**).
